# Direct M2 macrophage co-culture overrides viscoelastic hydrogel mechanics to promote fibroblast activation

**DOI:** 10.1101/2024.10.13.618034

**Authors:** Leilani R. Astrab, Mackenzie L. Skelton, Steven R. Caliari

## Abstract

Fibroblast activation drives fibrotic diseases such as pulmonary fibrosis. However, the complex interplay of how tissue mechanics and macrophage signals combine to influence fibroblast activation is not well understood. Here, we use hyaluronic acid hydrogels as a tunable cell culture system to mimic lung tissue stiffness and viscoelasticity. We applied this platform to investigate the influence of macrophage signaling on fibroblast activation. Fibroblasts cultured on stiff (50 kPa) hydrogels mimicking fibrotic tissue exhibit increased activation as measured by spreading as well as type I collagen and cadherin-11 expression compared to fibroblasts cultured on soft (1 kPa) viscoelastic hydrogels mimicking normal tissue. These trends were unchanged in fibroblasts cultured with macrophage-conditioned media. However, fibroblasts directly co-cultured with M2 macrophages show increased activation, even on soft viscoelastic hydrogels that normally suppress activation. Inhibition of interleukin 6 (IL6) signaling does not change activation in fibroblast-only cultures but ameliorates the pro-fibrotic effects of M2 macrophage co-culture. These results underscore the ability of direct M2 macrophage co-culture to override hydrogel viscoelasticity to promote fibroblast activation in an IL6-dependent manner. This work also highlights the utility of using hydrogels to deconstruct complex tissue microenvironments to better understand the interplay between microenvironmental mechanical and cellular cues.

## 1. Introduction

Cells are embedded within tissues in extracellular matrix (ECM), a structural framework that provides physical support and cell-directing signals in both physiological and pathological contexts, such as wound healing and disease progression^[1]^. Cells sense and respond to the ECM through mechanotransduction, a process by which mechanical stimuli are converted to biochemical signals^[2]^. In the event of an injury, cells such as macrophages and fibroblasts are activated to facilitate wound healing by clearing debris, secreting cytokines to regulate inflammation, and producing ECM^[3,4]^. However, when the wound healing process is dysregulated, the dynamic relationship between ECM cues and resident cells contributes to the progression of fibrotic diseases^[4–6]^. In pulmonary fibrosis, the persistence of macrophages and stiffening of ECM promote fibroblast activation to myofibroblasts^[5,7,8]^. These activated fibroblasts are collagen-producing and contractile cells, crucial for scar formation during wound healing. In pulmonary fibrosis, however, chronic fibroblast activation leads to excessive scarring around alveoli at the site of gas exchange^[9,10]^. This disruption to lung architecture increases tissue stiffness, impairing lung function and often leading to organ failure^[11–13]^. To develop more effective therapeutics, it is essential to better understand the mechanisms that sustain fibroblast activation.

Macrophages have been implicated as crucial players in fibrotic disease. Specifically, macrophages have been shown to activate fibroblasts by producing pro-fibrotic factors and to amplify the activation response by secreting cytokines that recruit additional fibroblasts and inflammatory cells^[8,14–16]^. However, macrophages are highly plastic, exhibiting a broad range of phenotypes, functions, and tissue-specific roles that result in various secretory profiles^[17,18]^. Depending on their phenotype, macrophages release inflammatory mediators such as interleukin 6 (IL6), secrete growth factors like transforming growth factor-beta (TGFβ), phagocytose pathogens and debris, and regulate ECM turnover through the release of matrix metalloproteinases (MMPs) and tissue inhibitors of metalloproteinases (TIMPs)^[14,19]^. While the naming convention of classically-activated “M1” and alternatively-activated “M2” macrophages does not fully capture the complexity and heterogeneity of macrophage behaviors *in vivo*, this paradigm offers an approximation of polarization states replicated *in vitro*^[20,21]^.

Studies have shown that macrophage-induced fibroblast activation is phenotype-dependent, with M2 macrophages often implicated in sustained fibrogenesis^[3,7,20,22–24]^. For instance, M2 macrophages were shown to promote activation in human dermal fibroblasts, seen by increased levels of type I collagen and TGFβ1 expression as well as expression of the myofibroblast marker alpha smooth muscle actin (ɑSMA) compared to fibroblast-only cultures or transwell co-cultures with M0 or M1 macrophages^[25]^. While macrophages can activate fibroblasts through secreted factors, they also promote activation through direct cell-cell contact involving transmembrane proteins like cadherins^[26]^. One study demonstrated the influence of cadherin-11 (CDH11) in mediating direct crosstalk between M2 macrophages and myofibroblasts, showing that myofibroblasts in these co-cultures exhibited increased contractility, ɑSMA stress fiber formation, and increased levels of TGFβ production compared to fibroblasts cultured alone or in segregated co-cultures^[20]^. CDH11 is a transmembrane protein involved in cell adhesion that is crucial for proper tissue development, however, recent evidence has demonstrated its upregulation in several fibrotic diseases, including pulmonary fibrosis^[20,27,28]^. Lung sections from patients with idiopathic pulmonary fibrosis exhibit elevated CDH11 expression compared to normal controls. Similarly, bleomycin-treated mice exhibit increased CDH11 levels, while CDH11-deficient mice exhibit less severe fibrosis when challenged with bleomycin^[20,28]^. Although these findings highlight the role of M2 macrophages and CDH11-mediated crosstalk in fibrosis progression, the complexity of the *in vivo* environment makes it challenging to isolate the role of any one factor on fibroblast activation. Furthermore, with macrophage crosstalk and changing ECM mechanics both playing key roles in driving fibrogenesis, there is a need to investigate their relative contributions in controlled *in vitro* microenvironments that recapitulate native tissue mechanics.

Utilizing hydrogels as *in vitro* models has proven valuable for facilitating the investigation of physiologically-relevant cell behaviors while limiting confounding variables arising *in vivo*^[29,30]^. These hydrophilic crosslinked polymer networks can be fabricated from a variety of synthetic or naturally-occurring materials, enabling control over salient extracellular environmental features such as mechanics. By utilizing orthogonal chemistries to incorporate different functional groups on polymer backbones, our group and others have been able to independently tune hydrogel mechanical properties like stiffness and viscoelasticity^[31–35]^. Tuning these characteristics enables the investigation of their respective influence on cell behavior and the development of hydrogels that mimic both normal and diseased tissue environments, providing insights that would be impossible to glean from traditional studies using tissue culture plastic^[31,36–38]^. This approach allows us to bridge the gap between complex *in vivo* models and standard *in vitro* substrates like plastic and glass that do not mimic tissue mechanics. Studies from our group have shown that increasing hydrogel stiffness and reducing viscoelasticity result in direct changes to cellular function and morphology^[32,33,36]^. Notably, fibroblasts seeded on stiff elastic substrates mimicking fibrotic tissue exhibit increased signs of activation like increased spread area, focal adhesion formation, and actin stress fiber organization^[33,36,37]^. Conversely, studies have shown that culturing cells on viscoelastic hydrogels mimicking normal tissue mechanics results in reduced spreading and decreased ɑSMA expression^[31,36]^. In the context of fibrotic disease, hydrogel systems can serve as platforms to better understand the underlying mechanisms driving pathological cell behavior, which could inform more effective therapeutic design.

While the combined role of macrophages and fibroblasts in fibrosis progression has been studied before, the signal integration of substrate mechanics and macrophage phenotype has not been widely explored. To decouple macrophage-fibroblast crosstalk from changing ECM mechanics, we utilize a hyaluronic acid (HA)-based hydrogel system to investigate macrophage-fibroblast co-cultures. Using a high-throughput, 96-well hydrogel array developed in our lab^[39]^, we demonstrate that this hydrogel platform supports the emergence of fibrotic cell behaviors seen in other *in vitro* and *in vivo* studies. This allows us to probe the influence of distinct macrophage phenotypes and modes of crosstalk on fibroblast activation in hydrogels with relevant normal and fibrotic mechanics. Additionally, with this system we can independently tune hydrogel stiffness and viscoelasticity on a per well basis, allowing us to test how these mechanical properties are individually contributing to macrophage-fibroblast signaling.

## 2. Results and Discussion

### 2.1 Hyaluronic acid-based hydrogels were designed with tunable stiffness and viscoelasticity

Previous studies from our group and others have demonstrated that fibroblasts exhibit signs of activation on stiffer (Young’s moduli >10 kPa) elastic substrates^[32,36,39]^. Additionally, there have been extensive studies demonstrating the influence of macrophage phenotype on fibroblast activation^[6,7,21]^. However, many of the systems used to investigate the relationship between fibroblasts and macrophages do not capture the mechanics of normal or fibrotic lung tissue and are relatively low-throughput, limiting the variables one can perturb in a single experiment. Here, we employ a 96-well-based hydrogel platform developed in our lab that addresses these challenges, allowing us to fabricate hydrogels with independently tunable storage and loss moduli^[39]^. With this system we can fabricate distinct hydrogel microenvironments in each well, allowing us to explore various mechanical influences and forms of cell crosstalk.

Leveraging the inherent biocompatibility and ease of modification of hyaluronic acid (HA), we incorporated norbornene groups to synthesize a modified polymer (NorHA) that reacts with free thiols via light-mediated thiol–ene click chemistry (**Fig. 1A**). This system allows for user-defined control over hydrogel storage modulus by varying the ratio of dithiol crosslinker to available norbornenes, thereby controlling the degree of covalent crosslinking. Additionally, we synthesized β-cyclodextrin-modified hyaluronic acid (CDHA) and a thiolated adamantane peptide to employ reversible guest-host interactions that allow us to tune hydrogel loss modulus, enabling control over hydrogel viscoelasticity^[36]^ (**Fig. 1A**). The adamantane peptide can click onto norbornene groups, where it’s free to physically interact with the hydrophobic cavity of β-cyclodextrin; however, under cell-relevant traction forces, these groups can disassemble to give the hydrogel network viscous character. These modified polymers enabled the fabrication of hydrogels with a range of Young’s moduli (1-50 kPa) that encompassed mechanical properties of normal and fibrotic lungs^[40,41]^.

**Figure 1.**
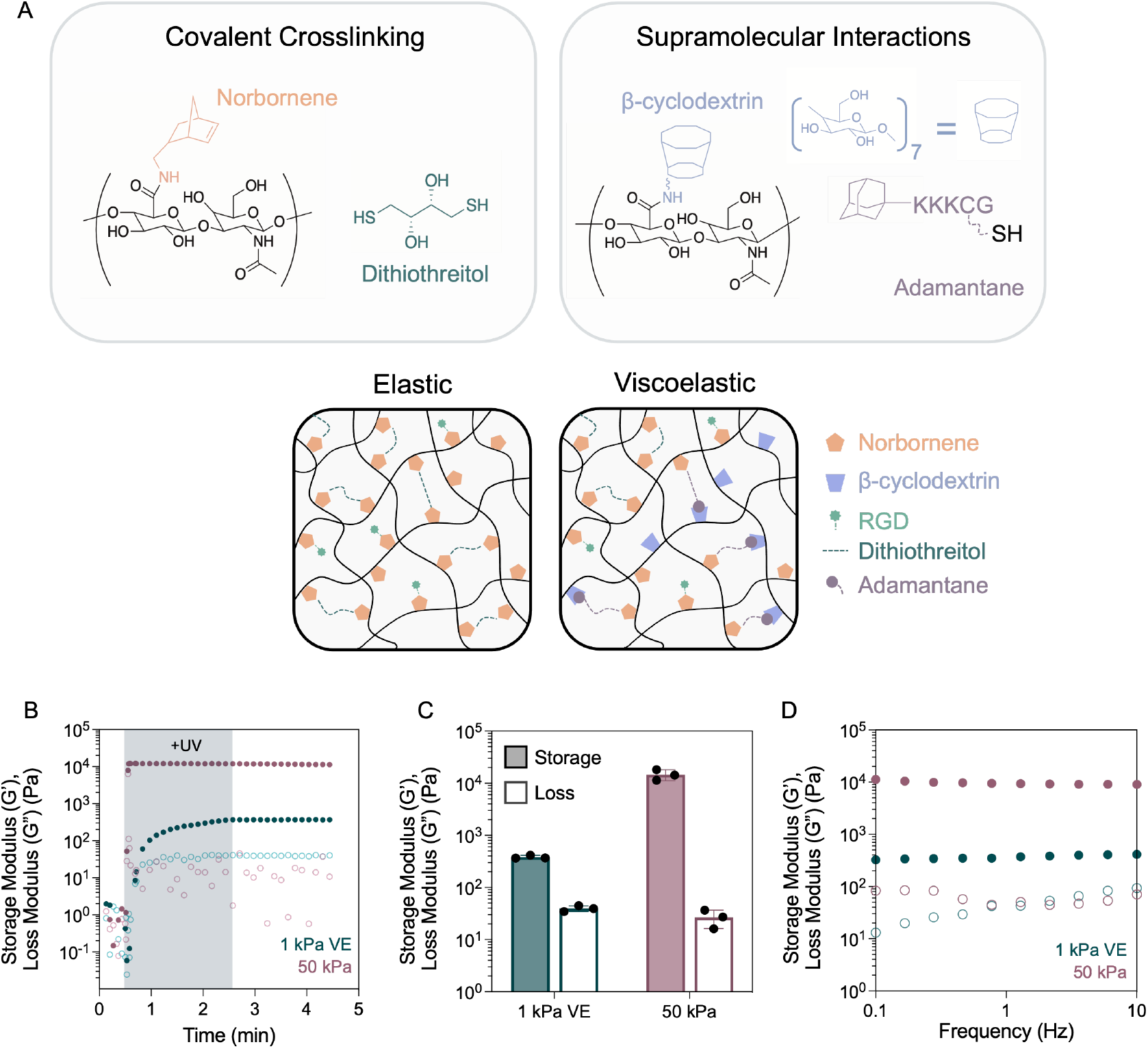
Design and characterization of a viscoelastic hydrogel platform to mimic tissue mechanics. A) Schematic of covalent (thiol-ene addition) and supramolecular (Ad-CD guest-host) crosslinking mechanisms used in this hydrogel system to tune hydrogel storage and loss moduli respectively. B) Characterization of soft, 1 kPa viscoelastic (blue) and stiff, 50 kPa elastic (purple) hydrogel mechanical properties using oscillatory shear rheology. Storage moduli (G’) are shown in closed circles and loss moduli (G”) are shown in open circles. Time sweeps (1 Hz, 1% strain) showing UV-mediated gelation (365 nm, 5 mW cm^-2^, 2 min) of elastic and viscoelastic hydrogel formulations. C) Average storage (G’) and loss (G”) moduli values from the last 30 s of time sweeps. D) Frequency sweeps (0.1-10 Hz) demonstrate frequency-dependent behavior of 1 kPa viscoelastic hydrogel formulation.

To mimic mechanics of normal lung tissue, we formulated a hydrogel with a Young’s modulus of ∼ 1 kPa that incorporated viscoelasticity through guest-host physical interactions. This formulation had a tan delta of ∼ 0.1, indicating the ratio of storage to loss modulus is within an order of magnitude, characteristic of normal soft tissues like lung^[11,41,42]^ (**Fig. 1B, C**). We modeled fibrotic lung tissue utilizing an elastic hydrogel with purely covalent crosslinks resulting in a Young’s modulus of ∼ 50 kPa (**Fig. 1B, C**). *In situ* rheology showed that hydrogel precursors with photoinitiator remained liquid until the application of UV light where gelation occurred rapidly, and moduli plateaued after 2 min. Hydrogel moduli remained stable after turning off the UV light (**Fig. 1B**). Oscillation frequency was varied to further characterize hydrogel viscoelasticity. Frequency sweep tests showed increasing hydrogel loss moduli with increasing frequency in the 1 kPa viscoelastic (VE) formulation but not the 50 kPa elastic hydrogel, indicating dissociation of adamantane and β-cyclodextrin guest-host interactions (**Fig. 1D**). Taken together, these results indicate the successful fabrication of hydrogels that mimic normal (soft, viscoelastic) and fibrotic (stiff, elastic) lung tissue.

### 2.2 Fibroblasts exhibit increased spreading and expression of type I collagen and cadherin-11 in response to stiff elastic substrates

After fabricating hydrogels that mimicked the stiffness and viscoelasticity of normal and fibrotic lung tissue, we next wanted to understand how fibroblasts would respond to differences in these mechanics. We utilized a 96-well hydrogel array developed in our lab to culture human lung fibroblasts on either 1 kPa VE or 50 kPa elastic hydrogels for 2 days (**Fig. 2**)^[39]^. Cells were then fixed and stained to evaluate metrics of fibroblast activation: cell spread area, cell circularity (measured by cell shape index), type I collagen expression, and CDH11 expression^[20,27]^. Activated fibroblasts typically show increased spread area, reduced circularity/cell shape index, and increased type I collagen expression. CDH11 was chosen as a marker of interest due to its involvement in several fibrotic diseases, including pulmonary fibrosis^[27,28]^, as well as its aforementioned role in fibroblast-macrophage crosstalk in fibrosis^[20]^. We found that fibroblasts exhibited significantly increased spread area and decreased circularity (shown by decreased cell shape index) when cultured on 50 kPa hydrogels (**Fig. 2B-C**). Additionally, fibroblasts on these stiffer, elastic hydrogels exhibited increased expression of type I collagen and CDH11 (**Fig. 2D-E**). These results demonstrate that fibroblasts cultured on 50 kPa elastic hydrogels exhibited significant morphological differences and increased expression of fibrotic markers from those cultured on 1 kPa VE substrates, indicating that stiff, elastic substrate mechanics are sufficient to activate fibroblasts.

**Figure 2.**
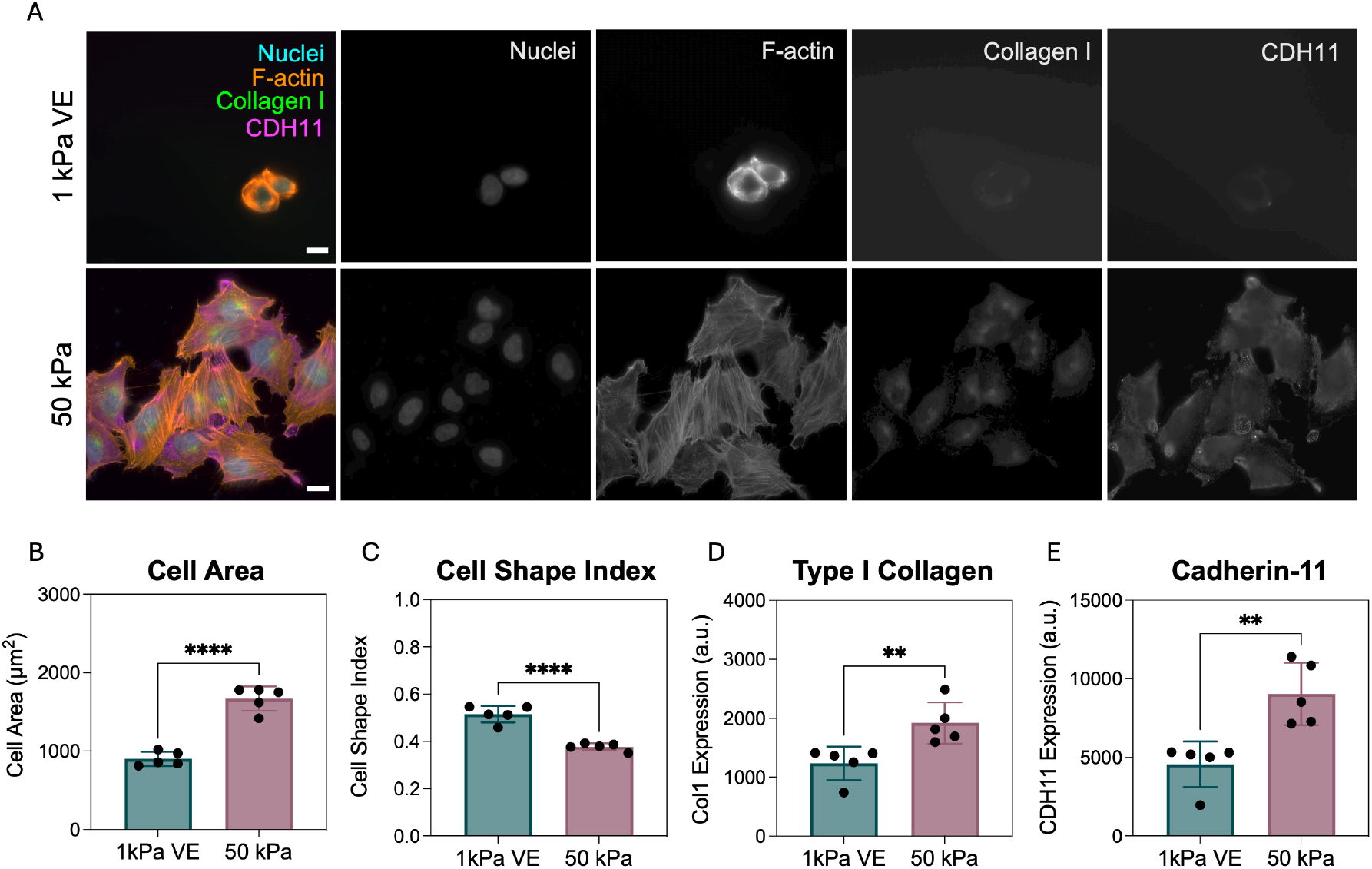
Fibroblasts seeded on stiff elastic hydrogels mimicking fibrotic lung mechanics exhibit increased activation compared with fibroblasts on hydrogels mimicking normal lung mechanics. A) Representative images of fibroblasts seeded on 1 kPa viscoelastic hydrogels (top row) and 50 kPa elastic hydrogels (bottom row) after 2 days of culture. Cells stained for nuclei (cyan), F-actin (orange), type I collagen (green), and cadherin-11 (magenta). Scale bars = 20 μm. Image quantification demonstrates fibroblasts on 50 kPa elastic hydrogels exhibit significantly B) increased cell area, C) decreased cell shape index, which correlates to decreased cell circularity, D) increased type I collagen expression, and E) increased cadherin-11 expression compared to fibroblasts on 1 kPa viscoelastic hydrogels. *N* = 5 hydrogels per group, 242-264 individual cells per group. Statistical analyses performed via student’s t-tests. ****: *P* < 0.0001, **: *P* < 0.01.

To more thoroughly understand the individual roles of hydrogel stiffness and viscoelasticity in fibroblast activation, we conducted an additional study using hydrogels that mimicked normal (1 kPa) and fibrotic (50 kPa) tissue stiffness, both with and without viscoelastic properties (**Fig. S1**). We found that 1 kPa elastic hydrogels supported increased cell spreading and decreased cell circularity compared to 1 kPa VE substrates, highlighting the importance of incorporating viscoelasticity into hydrogel design. However, fibroblasts cultured on 50 kPa VE substrates did not show significant differences in activation metrics compared to fibroblasts on 50 kPa elastic substrates. Given these results, subsequent experiments continued with 1 kPa VE and 50 kPa elastic hydrogels to mimic normal and fibrotic lung tissue.

### 2.3 Fibroblasts cultured with macrophage-conditioned media respond similarly to hydrogel mechanics regardless of macrophage phenotype

After establishing that fibroblasts cultured on substrates mimicking fibrotic tissue mechanics exhibit increased activation compared to those on hydrogels mimicking normal tissue, we next sought to use this system to explore the influence of macrophage signaling on fibroblast activation. Macrophages are known key regulators of fibroblast activation in pulmonary fibrosis progression^[3,4,7,43]^, however the interplay of macrophage signals and substrate mechanics, particularly viscoelasticity, has not been widely explored. To probe this, we cultured fibroblasts on 1 kPa VE or 50 kPa hydrogels and treated them with conditioned media from M0, M1, or M2 macrophage cultures. Macrophages were polarized to M1 and M2 phenotypes using well-established protocols outlined in **Figure 3** and detailed in the Experimental Section^[44]^. Following polarization, media conditioned by macrophages was added to fibroblasts seeded on either 1 kPa VE or 50 kPa hydrogels (**Fig. 4, S2**). This approach allowed investigation of the influence of macrophage-derived soluble signals on fibroblast activation using physiologically-relevant substrates, while eliminating the impact of macrophage-fibroblast juxtacrine and reciprocal paracrine signaling.

**Figure 3.**
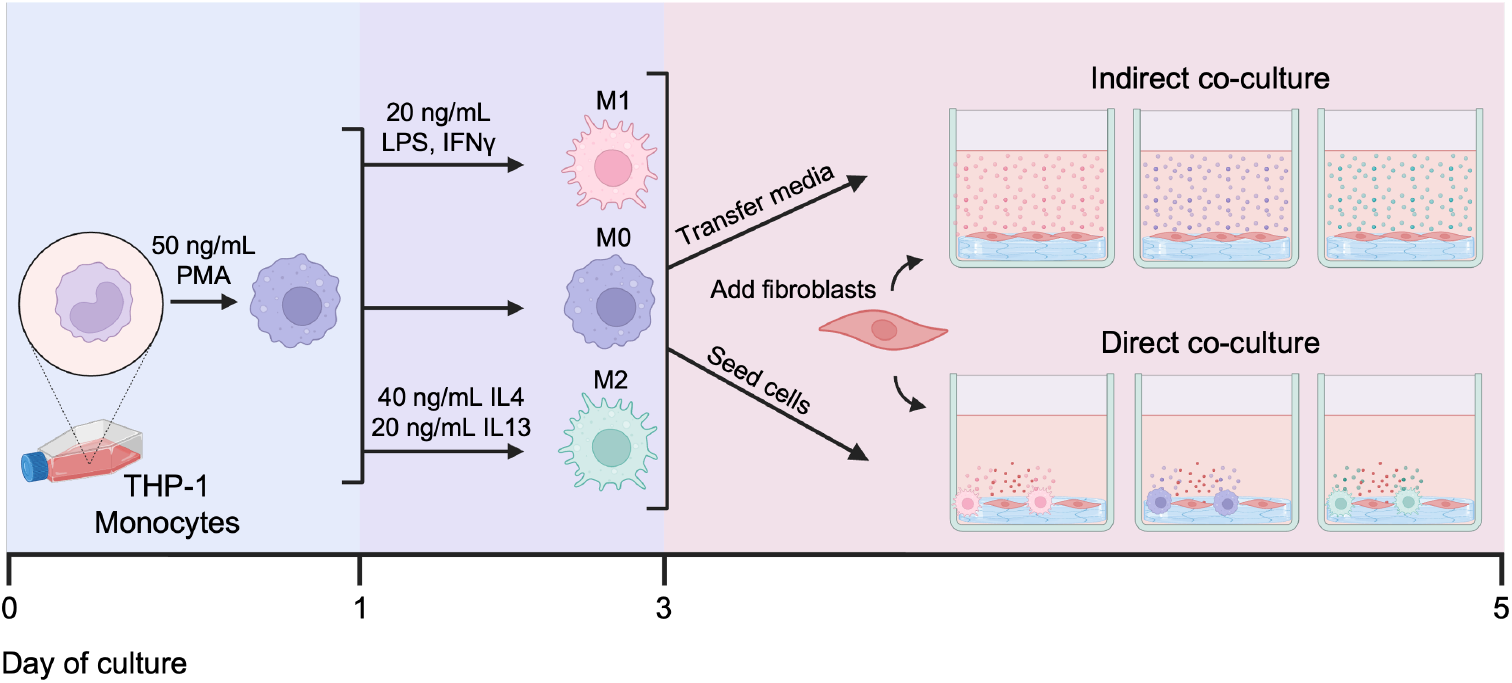
Timeline demonstrating cell culture workflow. Monocytes were differentiated to macrophages by adding 50 ng/mL of phorbol 12-myristate 13-acetate (PMA) to culture media for 1 day. Macrophages were then polarized to M0, M1, or M2 phenotypes by culturing in either base culture media, lipopolysaccharide (LPS, 20 ng/mL) and interferon gamma (IFNγ, 20 ng/mL), or interleukin 4 (IL4, 40 ng/mL) and interleukin 13 (IL13, 20 ng/mL), respectively. After 2 days, macrophage media was collected for conditioned media (indirect co-culture) experiments or macrophages were seeded with fibroblasts for direct co-culture studies. Co-cultures proceeded for 2 days before cell fixation for subsequent staining and analysis of fibroblast activation.

**Figure 4.**
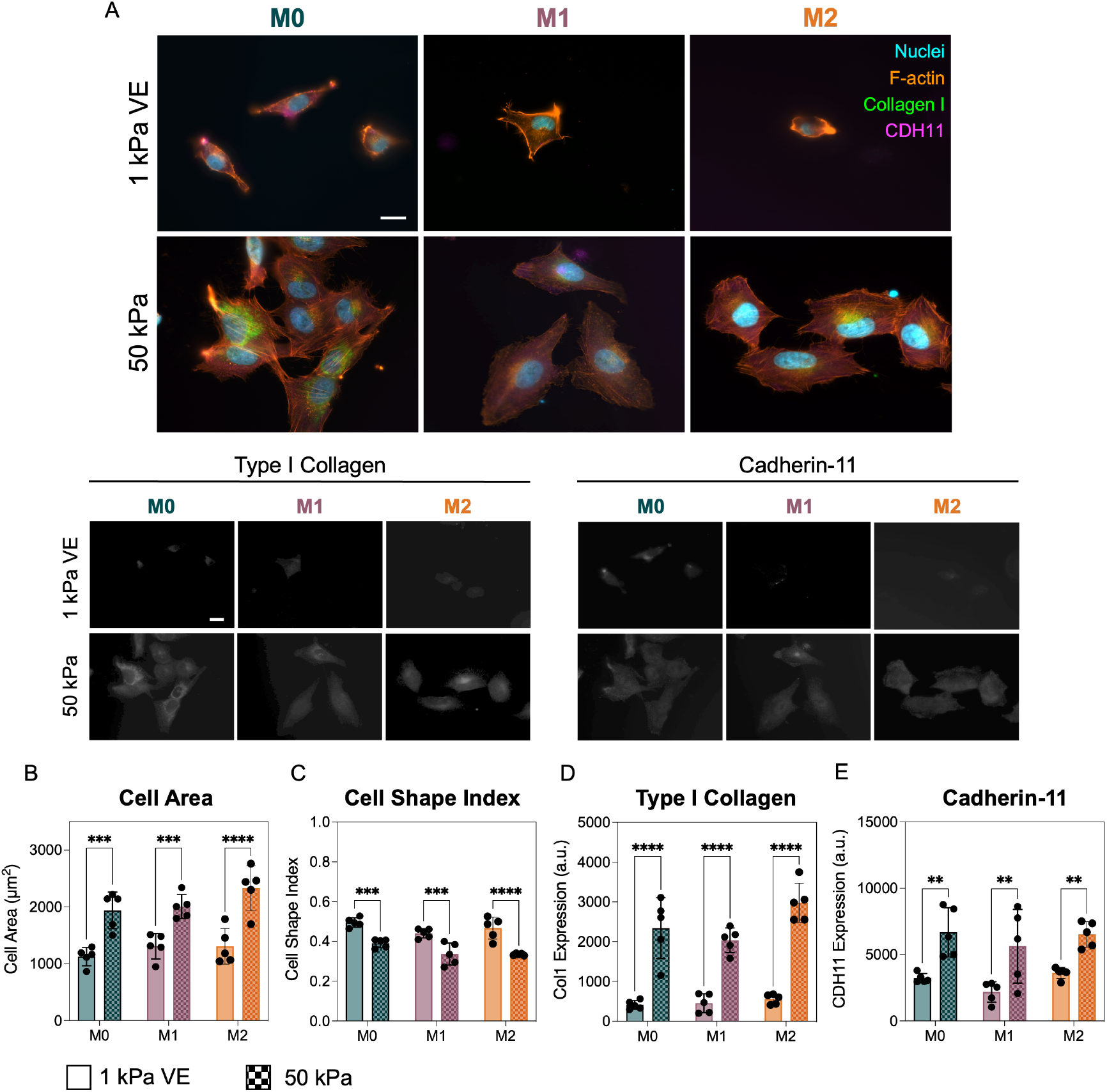
Fibroblasts cultured with conditioned media from M0, M1, or M2 macrophages seeded on hydrogels mimicking normal or fibrotic lung mechanics. A) Representative images of fibroblasts seeded on 1 kPa viscoelastic hydrogels (top row) and 50 kPa elastic hydrogels (bottom row) after 2 days of culture. Scale bars = 20 μm. Image quantification demonstrates fibroblasts on 50 kPa elastic hydrogels showed significantly B) increased spread area, C) decreased circularity measured by cell shape index, D) increased type I collagen expression, and E) increased cadherin-11 expression compared with fibroblasts on 1 kPa substrates regardless of macrophage-conditioned media type. *N* = 5 hydrogels per group, 124-243 individual cells per group. Statistical analyses performed via two-way ANOVA with Tukey’s HSD post-hoc testing. ****: *P* < 0.0001, ***: *P* < 0.001, **: *P* < 0.01.

After 2 days of culture, fibroblasts on 50 kPa hydrogels exhibited significantly increased cell spread area (**Fig. 4B**), decreased circularity (**Fig. 4C**), as well as increased type I collagen (**Fig. 4D**) and CDH11 (**Fig. 4E**) expression regardless of macrophage conditioned media type. These trends were consistent with what we observed with fibroblasts cultured in normal media. Furthermore, when compared to fibroblast-only data from a separate study, fibroblasts cultured in macrophage-conditioned media did not exhibit significant differences in any of these activation markers. This finding aligns with other studies, which reported that conditioned media from alveolar macrophages did not significantly influence the migration, proliferation, mRNA expression, or contractile activity of fibroblasts^[45,46]^. However, two studies found the opposite, demonstrating that fibroblasts treated with macrophage-conditioned media resulted in up-regulated inflammatory cytokine expression, expression of ECM-degrading enzymes, and increased proliferation^[47,48]^. The first study noted that fibroblasts treated with M1 macrophage-conditioned media showed increased expression of inflammatory cytokines but did not show up-regulation of *Acta2*, the gene encoding for ɑSMA^[47]^. The other study concluded that direct co-culture with macrophages elicited a fibroblast response more representative of *in vivo* characteristics than fibroblasts cultured with conditioned media^[48]^. Overall, our results suggest that soluble signals from macrophages alone are not sufficient to drive fibroblast activation on substrates with physiologically-relevant mechanical properties.

### 2.4 Fibroblasts directly co-cultured with M2 macrophages activate on soft viscoelastic hydrogels

After exploring the influence of macrophage soluble signals on fibroblast activation, we wanted to investigate the influence of direct crosstalk. We performed co-culture experiments where fibroblasts were directly cultured with M0, M1, or M2 macrophages on either 1 kPa VE or 50 kPa hydrogels for 2 days (**Fig. 5, S3**). We found that fibroblasts cultured on 50 kPa hydrogels directly with M0 or M1 macrophages exhibited increased spread area (**Fig. 5B**), decreased circularity (**Fig. 5C**), and increased expression of type I collagen (**Fig. 5D**) compared to 1 kPa VE hydrogels. In contrast, fibroblasts directly cultured with M2 macrophages exhibit these signs of activation on both 1 kPa VE and 50 kPa substrates, with no significant differences between hydrogel groups in any of these activation markers (**Fig. 5 B-D**). Additionally, fibroblast CDH11 expression did not increase on 50 kPa hydrogels in the direct M0 or M1 cultures compared to the 1 kPa VE hydrogels, unlike observations in the fibroblast-only and conditioned media experiments (**Fig. 5E**). However, there was a significant increase in CDH11 expression in fibroblasts co-cultured with M2 macrophages on both 1 kPa VE and 50 kPa hydrogels (**Fig. 5E**). Furthermore, we confirmed that fibroblast activation was consistent in these direct co-cultures with M2 macrophages by seeding cells on additional soft elastic and stiff viscoelastic substrates. We found that regardless of substrate stiffness or viscoelasticity, fibroblasts exhibited increased spread area, decreased circularity, and increased expression of type I collagen when compared to fibroblast-only controls (**Fig. S1**). These results demonstrate that direct culture with M2 macrophages leads to fibroblast activation, even on hydrogel substrates with compliant viscoelastic mechanics that would normally suppress activation. This supports data from other groups suggesting that macrophage-induced fibroblast activation is dependent on cell proximity^[20,45]^. Additionally, these data support emerging evidence highlighting the role CDH11 plays in fibrogenesis, specifically in mediating M2 macrophage-fibroblast crosstalk^[20,27]^.

**Figure 5.**
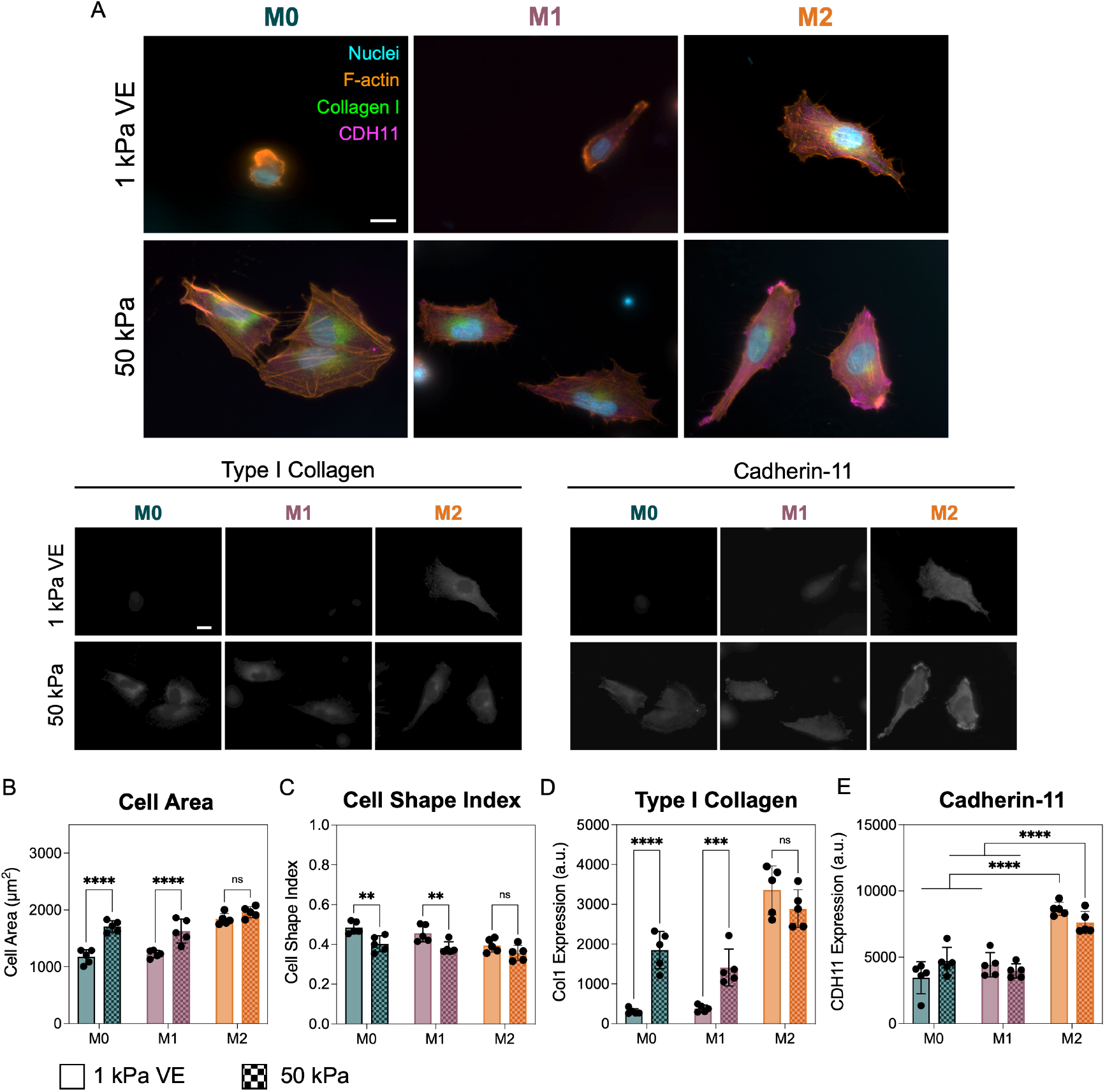
Fibroblasts directly co-cultured with M0, M1, or M2 macrophages seeded on hydrogels mimicking normal or fibrotic lung mechanics. A) Representative images of fibroblasts seeded on 1 kPa viscoelastic hydrogels (top row) and 50 kPa elastic hydrogels (bottom row) after 2 days of co-culture. Scale bars = 20 μm. B) Fibroblast spread area increased on 50 kPa hydrogels relative to 1 kPa VE hydrogels in M0 and M1 macrophage co-cultures, however M2 macrophage co-culture also led to increased fibroblast spreading on 1 kPa VE hydrogels. C) Cell circularity, measured by cell shape index, decreased in fibroblasts cultured on 50 kPa hydrogels relative to 1 kPa VE hydrogels in M0 and M1 macrophage co-cultures. However, fibroblasts co-cultured with M2 macrophages also displayed decreased circularity on 1 kPa VE hydrogels. D) Type I collagen expression increased on 50 kPa hydrogels relative to 1 kPa VE hydrogels in M0 and M1 macrophage co-cultures, but fibroblasts co-cultured with M2 macrophages exhibit increased type I collagen on 1 kPa VE hydrogels. E) Fibroblasts cultured with M2 macrophages exhibit increased levels of cadherin-11 relative to culture with M0 or M1 macrophages independent of hydrogel stiffness. *N* = 5 hydrogels per group, 165-278 individual cells per group. Statistical analyses performed via two-way ANOVA with Tukey’s HSD post-hoc testing. ****: *P* < 0.0001, ***: *P* < 0.001, **: *P* < 0.01.

To assess whether this co-culture-dependent fibroblast activation required direct physical contact with macrophages, we sorted images into two groups: those that had fibroblasts directly in contact with macrophages and those that did not. We then re-analyzed data with the same CellProfiler pipeline used for other experiments to investigate if there were significant changes in cell area, circularity, and type I collagen and CDH11 expression dependent on physical contact with macrophages. We found that fibroblasts in direct contact with M2 macrophages on 1 kPa VE substrates exhibited moderately increased spread area, type I collagen expression, and CDH11 expression as well as decreased circularity compared to those not in direct contact with macrophages (**Fig. S4**). However, even with these differences, fibroblasts not directly touching M2 macrophages still showed significantly increased spread area, type I collagen, and CDH11 expression compared with fibroblasts co-cultured with M0 or M1 macrophages (**Fig. S4**). While studies from other groups demonstrated that fibroblasts need to be in close proximity with M2 macrophages to induce fibroblast activation, some of these studies showed that distances of 100 μm (not accounting for cell height) are sufficient for fibroblast activation^[20,45]^. Additionally, previous work demonstrated that myofibroblasts can attract macrophages via force transmission through fibrillar matrices up to distances of 1300 μm^[49]^. In our work, the images analyzed to group touching and not touching cells had a field of view of 310 μm by 250 μm. Considering these results, it is likely that cells within this field of view, even if not seemingly physically touching, would be within the proximity resulting in increased fibroblast activation in previous studies^[20,45]^.

### 2.5 Interleukin 6 (IL6) inhibition diminishes fibroblast activation observed in direct M2 macrophage co-cultures

After observing that fibroblasts directly cultured with M2 macrophages exhibit increased signs of activation, even on soft viscoelastic hydrogels, compared to co-culture with M0 or M1 macrophages, we wanted to investigate how perturbing signaling pathways involved in fibroblast activation would impact this behavior. Fibroblasts were cultured alone or with M2 macrophages on 1 kPa VE and 50 kPa substrates. Actin polymerization, myosin II, Rho kinase signaling, TGFβ signaling, or IL6 signaling were inhibited through small molecule or antibody-based inhibitors outlined in **Table 1**. These targets were chosen for their established roles in mechanotransduction (actin, myosin II, Rho kinase signaling) or for their roles as cytokine signals implicated in fibroblast activation (TGFβ1, IL6)^[50–55]^.

**Table 1.** List of inhibitors used with type of inhibitor, final working concentration, and supplier information.

| Inhibitor | Inhibitor type | Target | Concentration | Supplier |
| --- | --- | --- | --- | --- |
| Anti-IL6R antibody [rhPM-1 (Tocilizumab)] | Antibody | Interleukin 6 receptor (IL6R) | 10 µg mL<sup>-1</sup> | Abcam (ab275982) |
| SB431542 | Small molecule | ALK5 receptor (TGFβ1R) | 50 µM | Abcam (ab120163) |
| Latrunculin A (LAT-A) | Small molecule | Actin polymerization | 5 µM | Abcam (ab144290) |
| (±)-Blebbistatin | Small molecule | Myosin II | 50 µM | Abcam (ab120425) |
| Y-27632 dihydrochloride | Small molecule | Rho kinase | 10 µM | Abcam (ab120129) |

Inhibition of actin polymerization led to decreased cell spread area, increased circularity, and decreased expression of type I collagen and CDH11 on all substrates compared to untreated controls (**Fig. 6A, S5**). This was expected as actin polymerization largely dictates cell shape metrics and plays a role in regulating collagen production^[56,57]^. Myosin II inhibition resulted in decreased cell circularity and decreased expression of type I collagen, however, there were no significant changes in cell area or CDH11 expression (**Fig. 6B, S6**). This was anticipated as myosin II is also responsible for cell morphology and synthesis of type I collagen^[58,59]^. However, these trends applied to all groups and therefore did not explain the unique activation seen with M2 macrophage-fibroblast co-cultures on compliant viscoelastic hydrogels. Inhibiting Rho kinase signaling led to decreased fibroblast circularity and type I collagen expression for 50 kPa hydrogel groups (**Fig. 6C, S7**). Additionally, Rho kinase inhibition decreased CDH11 expression in fibroblasts cultured with M2 macrophages. However, there was no significant change in CDH11 expression in fibroblast-only cultures. Previous studies showed that Rho kinase inhibition reduced CDH11 expression in mesenchymal stem cells, and that blocking CDH11 prevented activation of ROCK signaling^[60]^. Additionally, bleomycin-treated mice exhibited resolution of fibrosis when treated with a ROCK inhibitor, however this resolution did not occur when mice were depleted of macrophages before ROCK inhibition^[61]^. These findings suggest that macrophages play a crucial role in ROCK signaling during fibrosis progression, and that CDH11 may be involved in mediating this crosstalk.

**Figure 6.**
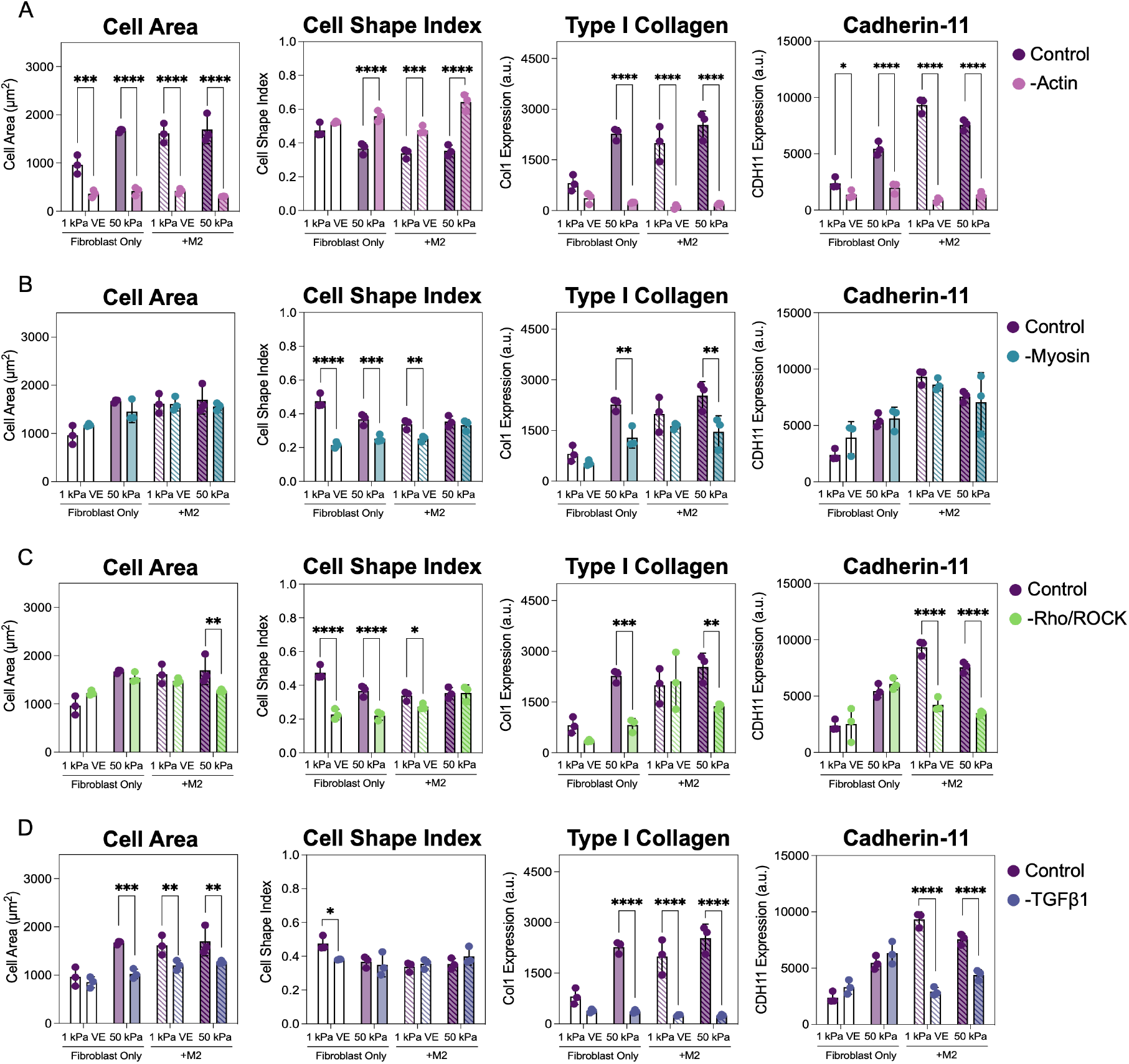
Investigating the influence of common mechanotransduction pathways and cytokine signals involved in fibroblast activation. Fibroblasts were cultured alone or with M2 macrophages on 1 kPa viscoelastic or 50 kPa elastic hydrogels for 2 days. A) Inhibiting actin polymerization resulted in decreased fibroblast spreading, more circular cells as indicated by increasing cell shape index, decreased type I collagen expression, and decreased cadherin-11 expression. B) Myosin II inhibition decreased cell circularity and type I collagen expression in fibroblasts on 50 kPa hydrogels. C) Inhibition of Rho kinase signaling decreased cell circularity, type I collagen expression in 50 kPa groups, and CDH11 expression in co-culture groups. D) Inhibition of TGFβ1 signaling decreased fibroblast spread area, type I collagen expression, and CDH11 expression in co-culture groups. *N* = 3 hydrogels per group, 58-180 individual cells per group. Statistical analyses performed via two-way ANOVA with Tukey’s HSD post-hoc testing. ****: *P* < 0.0001, ***: *P* < 0.001, **: *P* < 0.01, *: *P* < 0.05.

TGFβ1 is a known potent activator of fibroblasts, promoting synthesis of ECM proteins like collagen, and consequently driving scar tissue accumulation in fibrosis^[62]^. Blocking TGFβ1 signaling resulted in decreased fibroblast area on 50 kPa substrates in monocultures and in all co-culture groups compared to untreated controls (**Fig. 6D, S8**). Unsurprisingly, type I collagen expression was significantly decreased in all groups^[20,63]^. There was also significant reductions in CDH11 expression in fibroblast-M2 macrophage co-cultures but no change in fibroblast-only groups. Reduction of CDH11 in these co-cultures supports data showing that CDH11-mediated direct contact between fibroblasts and M2 macrophages establishes a compartment of active TGFβ1, thereby promoting fibroblast activation in these cultures^[20]^. In summary, inhibition of actin polymerization, myosin II, Rho kinase signaling, and TGFβ1 signaling resulted in changes to fibroblast morphology and expression of type I collagen and CDH11. However, perturbing these pathways did not reduce fibroblast activation markers in macrophage co-cultures to the levels observed in fibroblast-only cultures, and in many cases also changed the trends in the fibroblast-only cultures, suggesting the influence of these pathways was not specific to the M2 macrophage co-culture.

In fibroblast-M2 macrophage co-cultures, inhibition of IL6 impacted all four cell behavior metrics – cell area, circularity, and expression of type I collagen and CDH11 – regardless of hydrogel mechanics (**Fig. 7**). The changes in these metrics from blocking IL6 signaling resulted in fibroblast behavior comparable to the fibroblast-only controls on 1 kPa VE hydrogels. These controls promote a quiescent, non-activated fibroblast state, emphasizing the role of IL6 signaling in fibroblast activation. In contrast, fibroblast-only cultures did not exhibit any significant changes in activation markers following IL6 inhibition, emphasizing that this effect was unique to the interaction between fibroblasts and M2 macrophages. Pairwise statistical comparisons between all inhibitor groups are outlined in **Fig. S9**.

**Figure 7.**
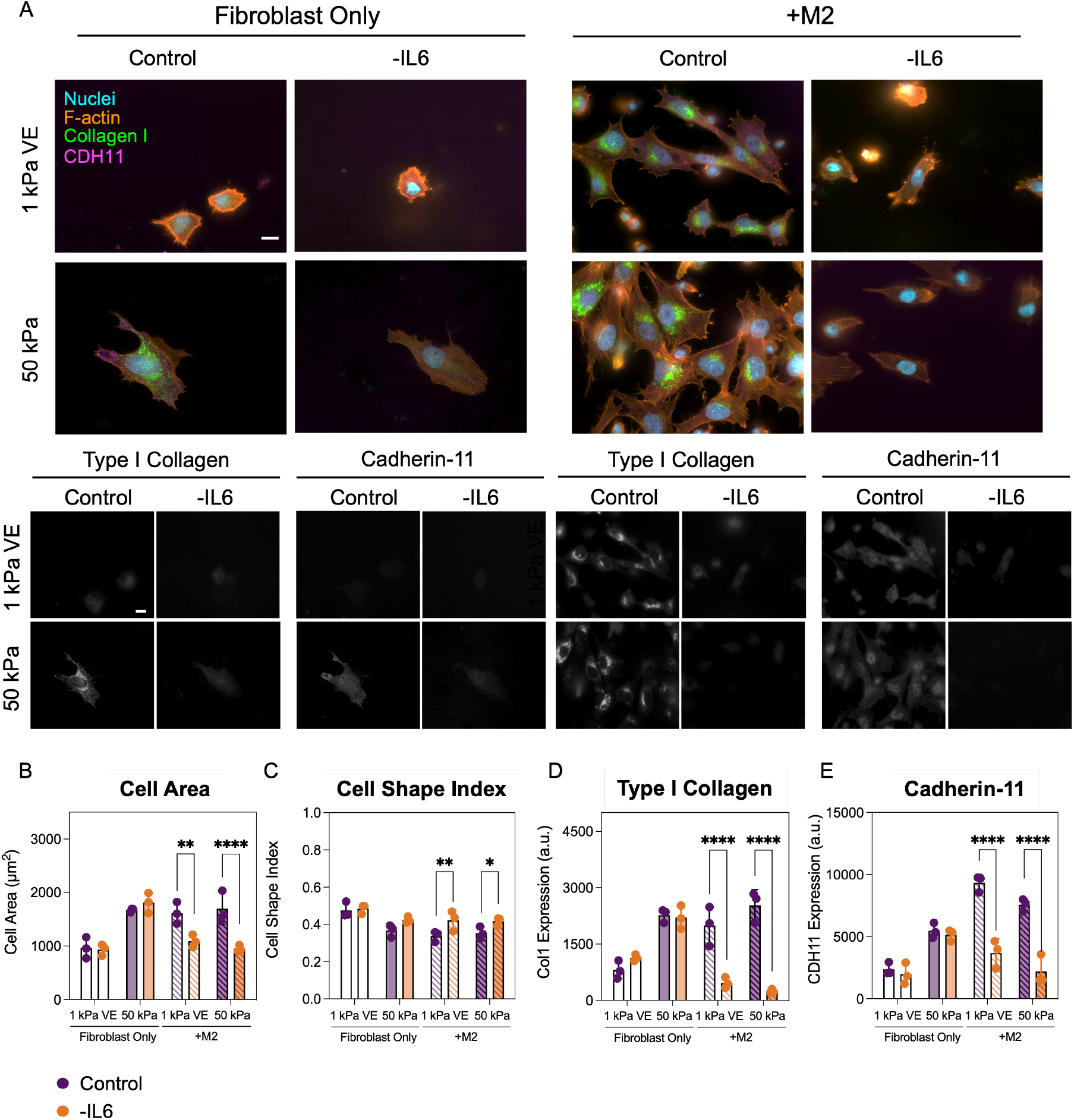
Interleukin 6 (IL6) signaling drives M2 macrophage-induced fibroblast activation on soft viscoelastic hydrogels. A) Representative images of fibroblasts cultured alone or with M2 macrophages on 1 kPa VE (top row) or 50 kPa (bottom row) hydrogels with or without an IL6 inhibitor. Scale bars = 20 μm. Blocking IL6 signaling B) reduces fibroblast spreading, C) increases circularity as measured by cell shape index, D) decreases type I collagen expression, and E) decreases cadherin-11 expression in fibroblasts cultured with M2 macrophages. Notably, IL6 inhibition brings fibroblast activation metrics back to the levels seen on 1 kPa VE control hydrogels without affecting activation metrics in fibroblast-only cultures. *N* = 3 hydrogels per group, 138-166 individual cells per group. Statistical analyses performed via two-way ANOVA with Tukey’s HSD post-hoc testing. ****: *P* < 0.0001, **: *P* < 0.01, *: *P* < 0.05. Images shown from the no inhibitor control groups are reproduced for all inhibitor experiment figures (Fig. 7, S5-S8).

In normal wound healing, IL6 is produced in early stages to stimulate immune cell chemotaxis, facilitate removal of necrotic tissue and debris, and promote production of other cytokines and growth factors^[64]^. After lung injury, IL6 is thought to originate from alveolar epithelial cells, then continues to be produced by macrophages and fibroblasts to promote ECM production in wound healing^[65]^. In bleomycin-induced lung fibrosis, IL6 functions as both a pro-inflammatory and pro-fibrotic factor, playing a role in early stages by protecting alveolar epithelial cells from reactive oxygen species and pneumocytes from bleomycin-induced apoptosis, while later promoting pro-fibrotic phenotypes of fibroblasts and macrophages^[66,67]^. The pleiotropic influence of IL6 was further demonstrated by studies showing IL6 neutralization could exacerbate or ameliorate bleomycin-induced lung fibrosis depending on the timing of neutralization. Specifically, neutralizing IL6 during the early inflammatory phase accelerated fibrosis, while neutralizing IL6 during the early fibrotic stage significantly reduced fibrosis^[68]^. These different effects could in part be due to the persistence of M2 macrophages in fibrosis progression, as studies have shown that IL6 enhances M2 macrophage polarization when combined with IL4 and IL13^[69]^. The addition of IL6 to these traditional M2 polarizing cytokines led to an increase in CD206 and arginase-1 positive cells, elevated arginase activity, and an increase in IL4 receptor-positive cells in bone marrow-derived macrophages compared to those stimulated with IL4 and IL13 alone^[69]^. This highlights IL6’s role in influencing macrophage-fibroblast crosstalk, where it may further amplify macrophage-driven fibroblast activation, contributing to fibrosis progression.

Given these inhibitor results and what others have shown in the field it seems that IL6 is an important soluble factor in mediating fibroblast activation in macrophage-fibroblast crosstalk. Yet, in our indirect co-culture studies, we did not observe the same effect of M2 macrophage-conditioned media on fibroblast activation. Thus, we propose several factors that may explain this. First, IL6 has a relatively short half-life of only a few hours, where our cultures were maintained for 2 days, meaning there may not have been sustained levels of IL6 required for fibroblast activation^[70,71]^. Additionally, IL6 inhibition did not reduce fibroblast activation in fibroblast-only cultures, indicating that IL6 plays a key role in promoting M2 macrophage-mediated fibroblast activation. This may be due to the enhancing effect IL6 has on M2 polarization. However, the complex signaling paradigm between these cells has not been fully elucidated, so it is likely that additional paracrine and/or juxtacrine signaling contribute to our observed results. Taken together, these findings highlight the importance of IL6 in mediating M2 macrophage-supported fibrogenesis.

## 3. Conclusions

In this study, we investigated the combined influence of macrophage phenotype and hydrogel mechanics on fibroblast activation. We utilized a high-throughput, hyaluronic acid-based hydrogel system to leverage thiol-ene chemistry and guest-host supramolecular interactions that allowed for control over hydrogel stiffness and viscoelasticity^[36,39]^. Using these hydrogels to model normal and fibrotic lung tissue, we found that fibroblasts show increased activation on 50 kPa elastic hydrogels mimicking fibrotic lung, demonstrated by increased spreading, decreased circularity, and increased type I collagen and cadherin-11 expression (**Fig. 8A**). These trends were not significantly changed in fibroblasts cultured with conditioned media from M0, M1, and M2 macrophages, indicating that macrophage soluble signals alone were insufficient to activate fibroblasts. However, we observed that direct co-culture with M2 macrophages overrides hydrogel mechanical signals, resulting in fibroblast activation even on a 1 kPa viscoelastic hydrogel that suppressed fibroblast activation in fibroblast-only and indirect co-culture experiments (**Fig. 8B**). We further identified that inhibiting IL6 signaling nullified activation driven by direct M2 macrophage crosstalk, returning fibroblasts to a quiescent state seen in fibroblast-only controls on 1 kPa VE hydrogels (**Fig. 8C**). Overall, these results demonstrate that direct culture with M2 macrophages can override the influence of substrate mechanics to activate fibroblasts, and that IL6 signaling plays a crucial role in mediating this response.

**Figure 8.**
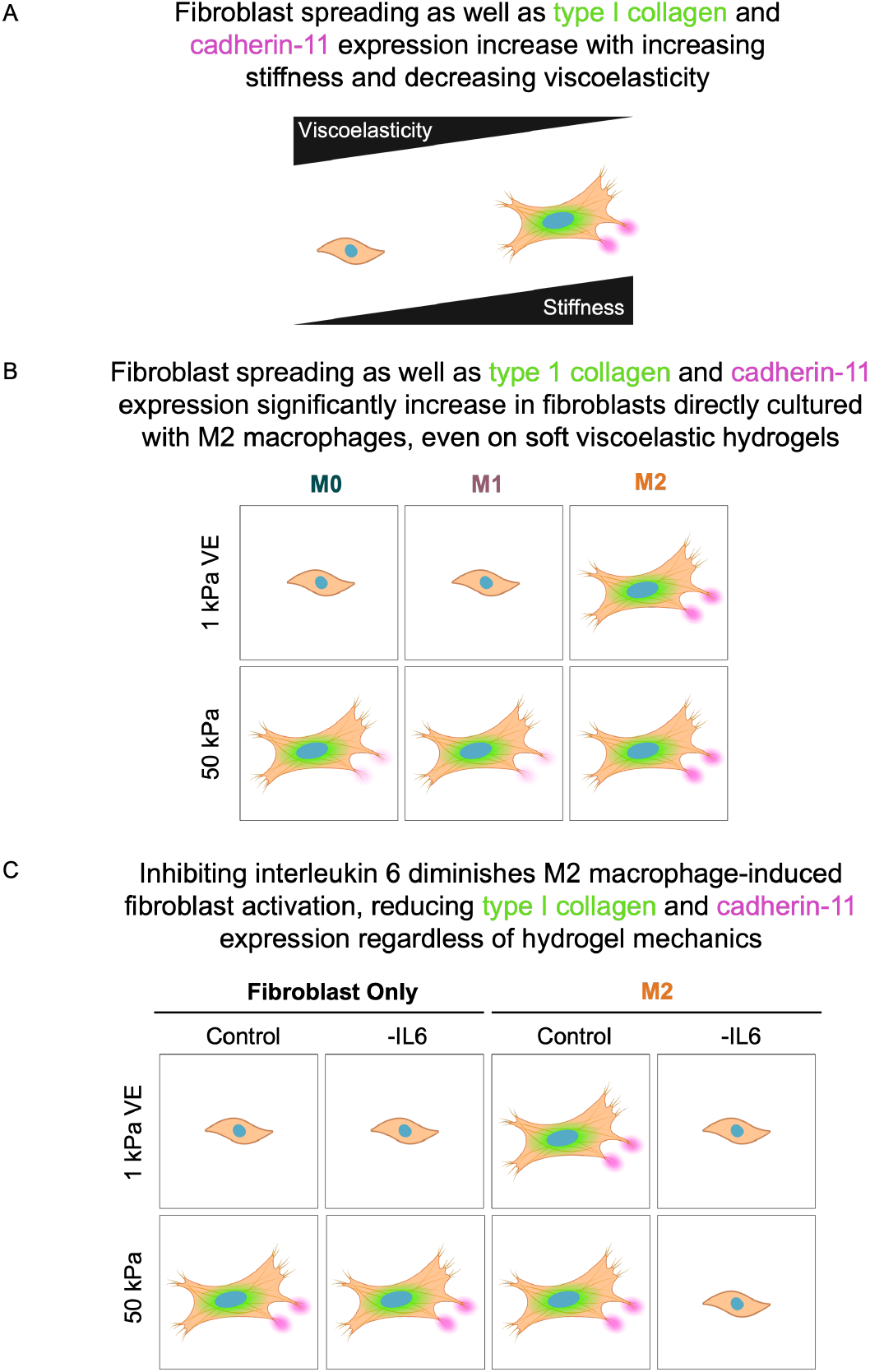
Summary of relationships between hydrogel mechanics, M2 macrophage co-culture, and interleukin 6 signaling on fibroblast activation. A) Fibroblasts exhibit increased spreading as well as type I collagen and cadherin-11 expression on stiffer, elastic substrates. B) Direct co-culture with M2 macrophages overrides mechanical cues to activate fibroblasts on 1 kPa VE hydrogels that mimic normal lung tissue mechanics. This is demonstrated by increased fibroblast spreading as well as increased expression of type I collagen and cadherin-11. C) Blocking interleukin 6 signaling nullifies M2 macrophage-induced fibroblast activation, reducing fibroblast spreading as well as type I collagen and cadherin-11 expression regardless of hydrogel mechanics.

## 4. Experimental Section

### 4.1 NorHA synthesis

Norbornene-modified HA was synthesized according to previously developed methods^[72]^. Briefly, sodium hyaluronate (Lifecore, 82 kDa) was reacted with Dowex 50W proton-exchange resin to form hyaluronic acid tetrabutyl ammonium salt (HA-TBA). This reaction solution was filtered, adjusted to a pH of 7.05, frozen, lyophilized, and the product confirmed using ^1^H NMR (500 MHz Varian Inova 500) (**Fig. S10**). HA-TBA was then reacted with 5-norbornene-2-methylamine and benzotriazole-1-yloxytris-(dimethylamino)phosphonium hexafluorophosphate (BOP) in dimethyl sulfoxide (DMSO) for 2 h at 25°C. The reaction was quenched with cold water, dialyzed (molecular weight cutoff 6-8 kDa) for 5 days, filtered to remove side products, dialyzed an additional 5 days, frozen, and lyophilized. The degree of modification was determined to be 29% by ^1^H NMR (**Fig. S11**).

### 4.2 β-CD-HDA synthesis

β-cyclodextrin hexamethylene diamine (β-CD-HDA) was synthesized using a previously outlined method^[73]^. Briefly, *p*-Toluenesulfonyl chloride (TosCl) dissolved in acetonitrile was added dropwise to a solution of β-cyclodextrin (CD) (5:4 molar ratio of TosCl:CD) at 25°C and allowed to react for 2 h. The reaction was then cooled on ice and an aqueous NaOH solution added dropwise (3.1:1 molar ratio of NaOH to CD). The reaction proceeded for 30 min at 25°C followed by addition of ammonium chloride to reach a pH of 8.5. The solution was cooled on ice, precipitated using cold water and acetone, and dried overnight. The CD-Tos product was then charged with hexamethylene diamine (HDA) (4 g/g CD-Tos) and dimethylformamide (DMF) (5 mL/g CD-Tos), then reacted under nitrogen at 80°C for 12 h. The reaction solution was then precipitated in cold acetone (5 × 50 mL acetone/1 g CD-Tos), washed with cold diethyl ether (3 × 100 mL), and dried. The β-CD-HDA product was confirmed using ^1^H NMR (**Fig. S12**).

### 4.3 CDHA synthesis

β-cyclodextrin-modified hyaluronic acid (CDHA) was synthesized via BOP-mediated coupling of β-CD-HDA with HA-TBA. The reaction was allowed to proceed for 3 h at 25°C in anhydrous DMSO, then quenched with cold water. The reaction solution was then dialyzed (molecular weight cutoff 6-8 kDa) for 5 days, filtered, dialyzed an additional 5 days, frozen, and lyophilized. The degree of modification was determined to be 25% by ^1^H NMR (**Fig. S13**).

### 4.4 HA hydrogel fabrication

To enable high throughput hydrogel fabrication, a 96-well plate array previously developed in our lab was utilized^[39]^. Glass pieces laser-cut to the dimensions of a bottomless 96-well plate (SI Howard Glass) were thiolated using (3-mercaptopropyl) trimethoxysilane, allowing covalent binding of norbornene groups within hydrogel solutions to the bottom of the well plate. Hydrogels were formed through ultraviolet (UV)-mediated thiol-ene reactions similar to previously reported methods^[36]^, and a bottomless 96 well plate was applied. Stiff elastic (50 kPa) NorHA hydrogel precursor solutions (5 wt% NorHA) containing 1 mM thiolated RGD peptide (GCGYGRGDSPG, Genscript) and dithiothreitol (DTT, thiol:norbornene ratio of 0.75) were photopolymerized (365 nm, 5 mW cm^-2^) in the presence of 1 mM lithium acylphosphinate (LAP) photoinitiator for 2 min. Soft viscoelastic (1 kPa VE) NorHA-CDHA hydrogels (3 wt% NorHA-CDHA) were fabricated by first mixing CDHA with a thiolated adamantane peptide (GCKKK-adamantane, Genscript) (1.2:1 molar ratio of Ad:CD) to first incorporate Ad–CD guest–host interactions. NorHA was next added to the precursor solution, along with DTT (thiol:norbornene ratio of 0.1), and RGD. The soft viscoelastic NorHA-CDHA solution was then photopolymerized using the same conditions as the stiff elastic group. Hydrogels swelled in phosphate-buffered saline (PBS) overnight at 37°C prior to use.

### 4.5 Rheological characterization

Rheological measurements were performed on an Anton Paar MCR 302 rheometer using a cone-plate geometry (25 mm diameter, 0.5°, 25 μm gap). Hydrogel mechanical properties were characterized using oscillatory time sweeps (1 Hz, 1% strain) with a 2 min UV curing step (365 nm, 5 mW cm^-2^) and oscillatory frequency sweeps (0.01–10 Hz, 1% strain).

### 4.6 Cell culture

Human lung fibroblasts (hTERT T1015 cell line purchased from Applied Biological Materials Inc.) were used before passage 5 for all experiments. Fibroblasts were cultured in Dulbecco’s modified Eagle medium (DMEM) supplemented with 10 v/v% fetal bovine serum (Gibco) and 1 v/v% penicillin/streptomycin/amphotericin B (1000 U mL^1^, 1000 μg mL^1^, and 0.25 μg mL^1^ final concentrations, respectively, Gibco). Cells were seeded at a concentration of 250,000 cells per 75 cm^2^ tissue culture flask and passaged every 3-4 days until ready for seeding on hydrogels. Prior to cell seeding, hydrogels were sterilized for at least 2 h by germicidal UV irradiation then incubated for at least 30 min in culture medium. Cells were then seeded into 96-well plate arrays (6 mm well diameter) at a density of 1.0 × 10^3^ fibroblasts per hydrogel and 3.3 × 10^5^ macrophages per hydrogel^[25,74]^. Cultures were allowed to proceed for 48 h before fixation and subsequent analysis.

THP-1 monocytes (TIB-202 cell line purchased from the American Type Culture Collection) were used between passages 6-8 for all experiments. Cells were maintained in RPMI 1640 (Gibco) medium supplemented with 10 v/v% heat-inactivated fetal bovine serum (FBS; Gibco), 1 v/v% penicillin/streptomycin/amphotericin B (1000 U mL^-1^, 1000 μg mL^-1^, and 0.25 μg mL^-1^ final concentrations, respectively, Gibco), 10 mM HEPES, 1 mM sodium pyruvate, 2.5 g L^-1^ D-glucose, and 0.05 mM β-mercaptoethanol. Cells were maintained between 0.25 and 0.5 × 10^6^ cells mL^-1^ with media replenishment every 3-4 days until ready for subsequent use. To differentiate monocytes to M0 macrophages, cells were seeded at 0.5 × 10^6^ cells mL^-1^ in 75 cm^2^ tissue culture flasks and stimulated with 50 ng mL^-1^ of PMA (phorbol 12-myristate 13-acetate; Sigma-Aldrich). After 24 h, cells were polarized to either M1 or M2 phenotypes through stimulation with lipopolysaccharide (LPS; 20 ng mL^-1^; Sigma Aldrich) and interferon gamma (IFN-γ; 20 ng mL^-1^; PeproTech) or interleukin 4 (IL4; 40 ng mL^-1^; PeproTech) and interleukin 13 (IL13; 20 ng mL^-1^; PeproTech), respectively^[44,75,76]^. After 48 h, cells were seeded onto hydrogels or prepped for flow cytometry. For indirect co-culture experiments, media was collected from M0, M1, or M2 macrophage cultures after polarization and added to fibroblasts that were seeded on hydrogels at a 1:1 ratio with fresh DMEM containing 10 v/v% heat-inactivated fetal bovine serum (FBS; Gibco) and 1 v/v% penicillin/streptomycin/amphotericin B (1000 U mL^-1^, 1000 μg mL^-1^, and 0.25 μg mL^-1^ final concentrations, respectively, Gibco). As with the direct co-culture experiments, cells were cultured for 48 h before fixing and proceeding with staining.

### 4.7 Immunocytochemistry, imaging, and analysis

Cells were fixed on hydrogels in 96-well plates with 10% neutral-buffered formalin for 15 min, then permeabilized using 0.1% Triton X-100 in PBS for 10 min. Bovine serum albumin (BSA; 3 w/v%) was used to block background staining for at least 2 h at room temperature. Hydrogels were then incubated overnight with primary antibodies against type I collagen (Col1, rabbit monoclonal anti-collagen I antibody, 1:200, Abcam ab138492) and cadherin-11 (CDH11), mouse monoclonal OB-cadherin/Cadherin-11 antibody 16A, 1:200, ThermoFisher MUB0306P). Hydrogels were washed 3 times with PBS then incubated for 2 h in the dark with rhodamine phalloidin to visualize F-actin (1:400) and secondary antibodies (AlexaFluor 488 goat anti-mouse IgG, 1:400; AlexaFluor 647 goat anti-rabbit, 1:400). After rinsing with PBS, hydrogels were then stained with DAPI nuclear stain (1:10000) for 1 min. Finally, hydrogels were rinsed twice with PBS and stored protected from light at 4°C until imaging. Microscopy was performed on a Zeiss AxioObserver 7 inverted microscope, with exposure time for each respective channel held constant throughout imaging.

A CellProfiler (Broad Institute, Harvard/MIT) pipeline was used to evaluate spread area, cell shape index (form factor), and staining intensity of Col1 and CDH11. Briefly, nuclei were identified using adaptive thresholding and overlaid with F-actin cytoskeletal staining to identify cells separately from background staining or debris. Cells with equal nuclear and cellular areas were deemed staining artifacts and eliminated from downstream analysis.

### 4.8 Inhibitor studies

Human lung fibroblasts and THP-1 monocyte-derived M2 macrophages were cultured as described above on either 1 kPa viscoelastic or 50 kPa elastic hydrogels. Wells were seeded with 6.0 × 10^3^ fibroblasts/hydrogel and 3.3 × 10^5^ M2 macrophages/hydrogel. Cells treated with antibody-based inhibitors were incubated on ice with antibody diluted in 2 v/v% FBS in PBS for 45 min, subsequently washed twice with 2 v/v% FBS in PBS, then resuspended in RPMI 1640 (Gibco) medium supplemented with 10 v/v% heat-inactivated fetal bovine serum (FBS; Gibco), 1 v/v% penicillin/streptomycin/amphotericin B (1000 U mL^-1^, 1000 μg mL^-1^, and 0.25 μg mL^-1^ final concentrations, respectively, Gibco), 10 mM HEPES, 1 mM sodium pyruvate, 2.5 g L^-1^ D-glucose, and 0.05 mM β-mercaptoethanol and seeded into the 96-well plate. Cells treated with small molecule inhibitors were left to adhere overnight before switching to inhibitor-containing media, while control wells and antibody-based treatment groups had media replaced with fresh supplemented RPMI. All groups were then cultured for an additional 24 h prior to fixation and subsequent staining as described above.

### 4.9 Statistical analysis

GraphPad Prism was utilized for all statistical analyses. Student’s t-tests (two experimental groups) or two-way ANOVA with Tukey’s HSD post-hoc tests (more than two experimental groups, independent variables: stiffness, macrophage co-culture) were performed. Each experiment involved a minimum of 3 hydrogels and/or 20 individual cells quantified per experimental group. Statistically significant differences are indicated by *, **, ***, **** corresponding to *P* < 0.05, 0.01, 0.001, or 0.001 respectively. Additional information regarding sample size or additional statistical analyses is in the figure captions.

## Supporting Information

Additional cell experiment results and ^1^H NMR spectra for the hydrogel components can be found in the Supporting Information.

## Acknowledgments

We thank Lily Burns for her assistance maintaining cells and synthesizing hydrogel materials used in this work, as well as Caliari Lab members for helpful discussions, support, and writing feedback. This work was supported by the NIH (R35GM138187) and NSF (GRFP to L.R.A. and M.L.S.). The content is solely the responsibility of the authors and does not necessarily represent the official views of the National Institutes of Health.

