## Supplemental Information for "Direct M2 macrophage co-culture overrides viscoelastic hydrogel mechanics to promote fibroblast activation"

S.R. Caliari
Department of Chemical Engineering
University of Virginia
Charlottesville, VA 22903, USA

keywords: hydrogels, fibrosis, macrophages, mechanotransduction

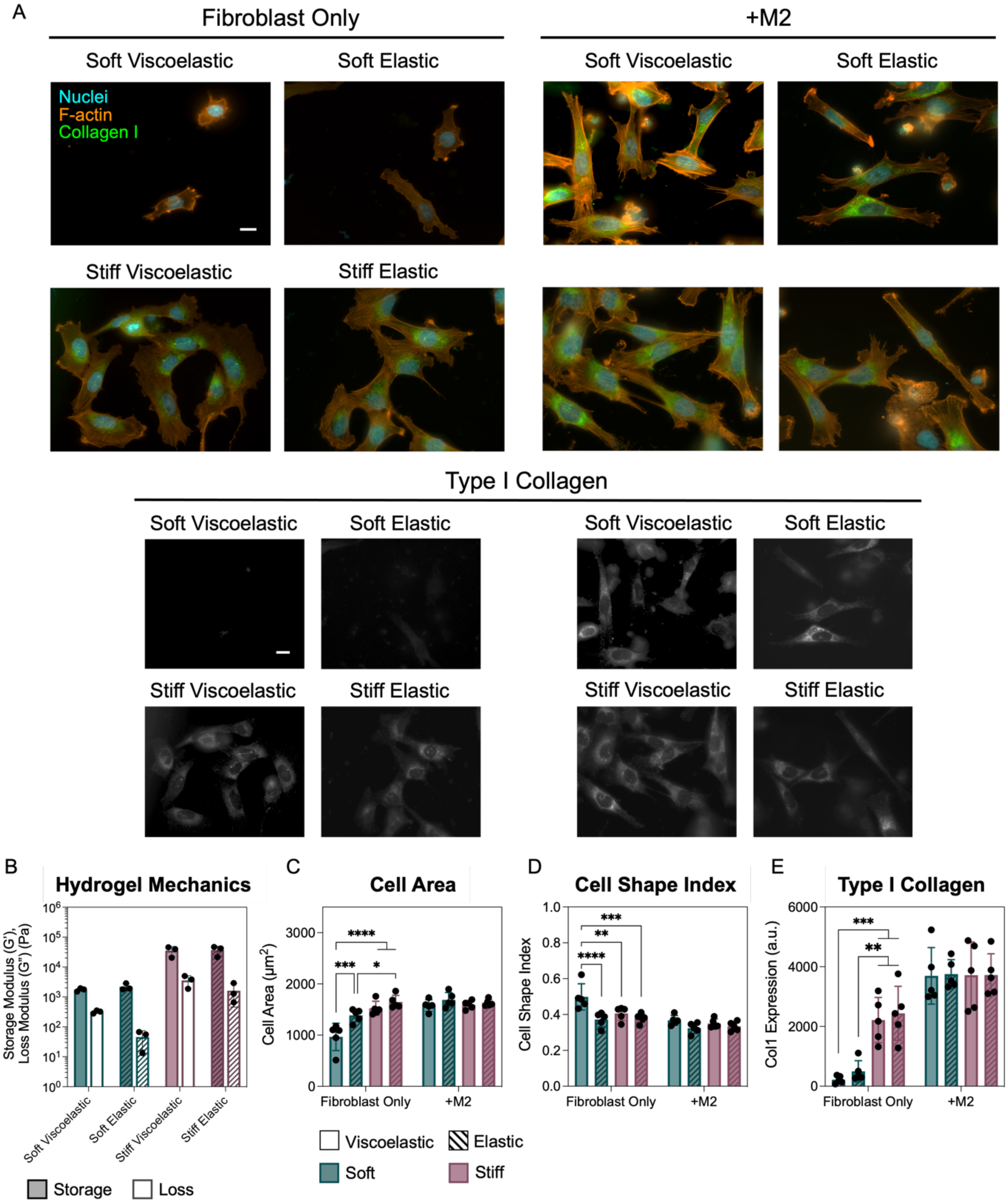

**Figure S1:** Fibroblasts cultured alone or with M2 macrophages on hydrogels of varying stiffness and viscoelasticity. A) Representative images of fibroblasts cultured on soft (1 kPa) viscoelastic, soft elastic, stiff (50 kPa) viscoelastic, and stiff elastic hydrogels after 2 days of culture. Scale bars = 20 μm. B) Average values of storage (G') and loss (G") moduli. Viscoelastic formulations exhibit loss moduli within an order of magnitude of storage moduli. C) Fibroblast spread area is significantly higher on all substrates compared to those on soft viscoelastic hydrogels in fibroblast-only cultures. No significant differences in fibroblast area observed in M2 macrophage co-cultures. D) Fibroblast circularity significantly decreased on all substrates compared to morphology on soft viscoelastic hydrogels in fibroblast-only cultures. No significant differences observed in M2 macrophage co-cultures. E) Fibroblast type I collagen expression significantly increased on stiff hydrogels in fibroblast-only cultures. No significant differences in expression observed in M2 macrophage co-cultures. $N = 5$ hydrogels per group, 126-444 cells per group. Statistical analyses performed via two-way ANOVA with Tukey's HSD post-hoc testing. ****: $P < 0.0001$, ***: $P < 0.001$, **: $P < 0.01$, *: $P < 0.05$.

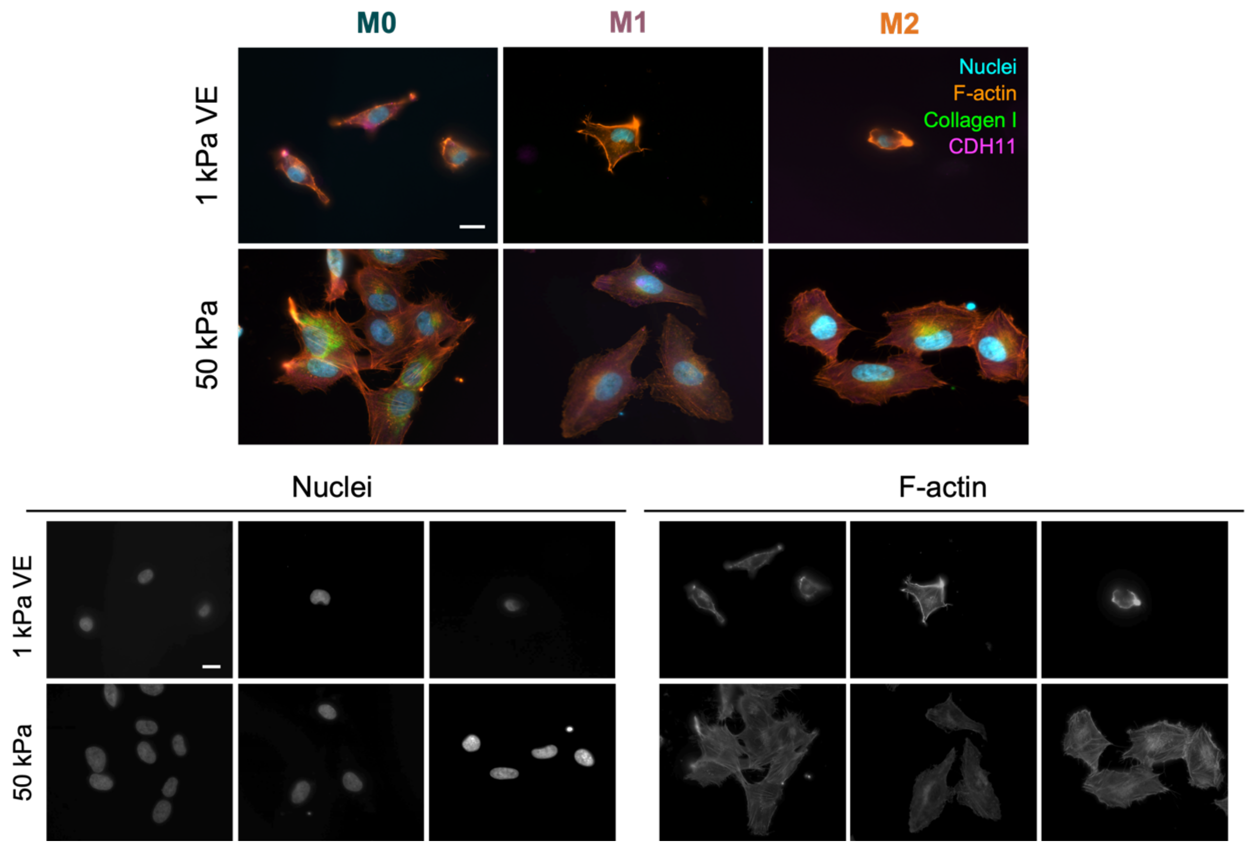

**Figure S2:** Representative and single channel images of nuclei and F-actin from fibroblasts cultured with M0, M1, or M2 macrophage-conditioned media from Figure 4. Scale bars = 20 μm.

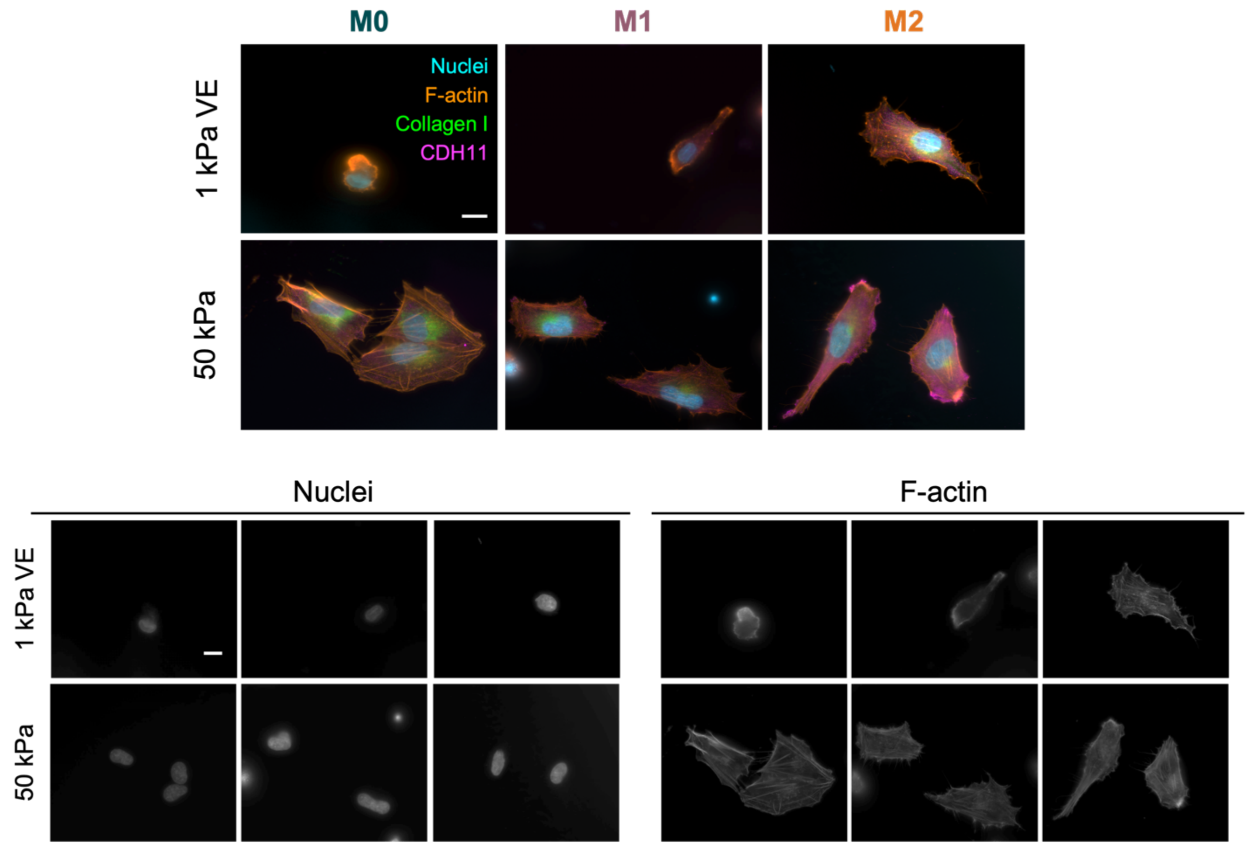

**Figure S3:** Representative and single channel images of nuclei and F-actin from fibroblasts directly cultured with M0, M1, or M2 macrophages from Figure 5. Scale bars = 20 μm.

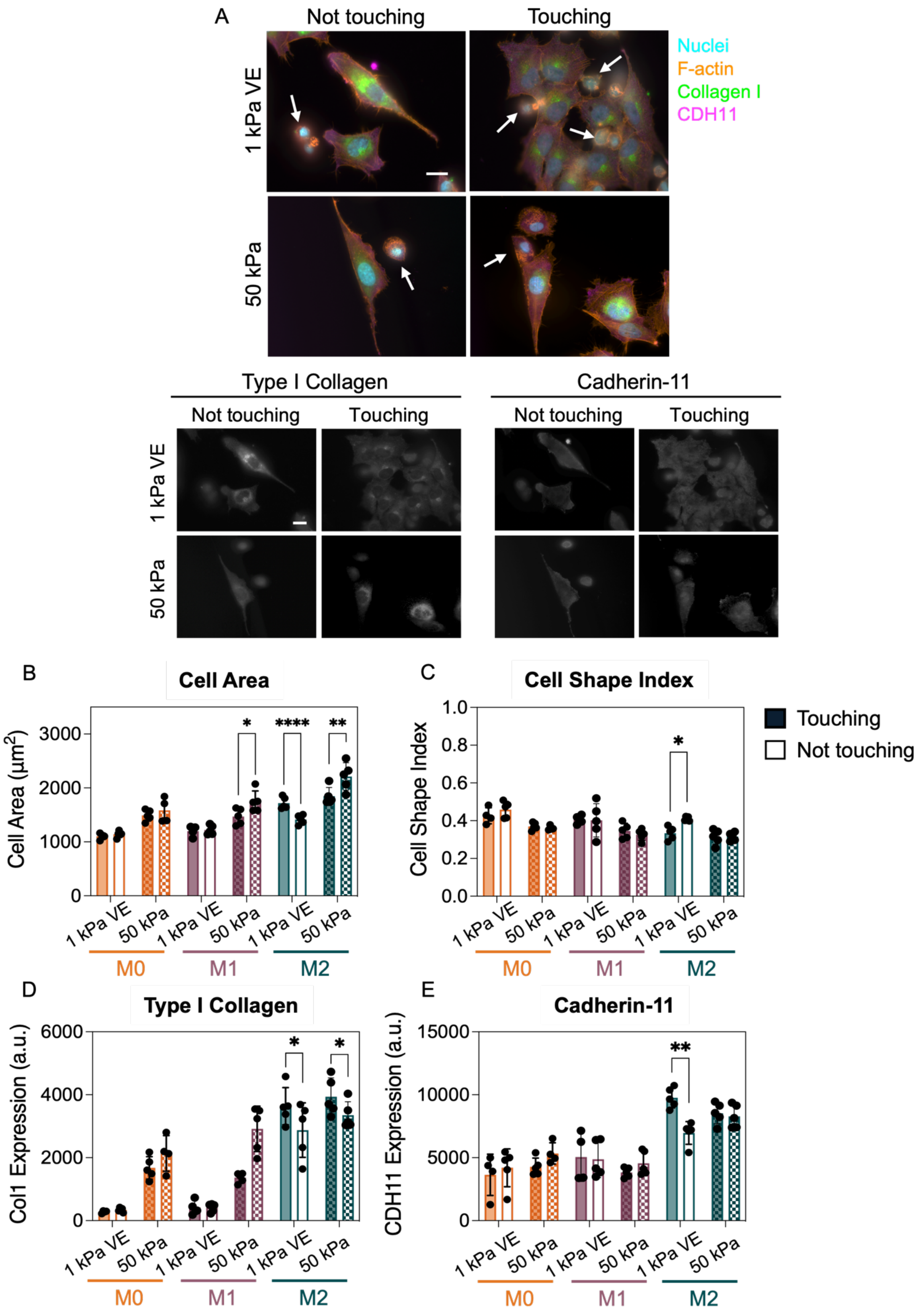
A
Not touching
Touching
Nuclei
F-actin
Collagen I
CDH11
1 kPa VE
50 kPa
Type I Collagen
Cadherin-11
B
Cell Area
Cell Area (μm2)
C
Cell Shape Index
Touching
Not touching
D
Type I Collagen
Col1 Expression (a.u.)
E
Cadherin-11
CDH11 Expression (a.u.)
M0
M1
M2

**Figure S4:** Fibroblast activation is not dependent on direct physical contact with M2 macrophages. Fibroblast-macrophage direct co-culture images were binned into two groups (touching and not touching), then re-analyzed to probe contact-dependent fibroblast activation. A) Representative images of fibroblasts touching and not touching M2 macrophages. White arrows indicate macrophages in images. Scale bars = 20 μm. Quantification of fibroblast B) spread area, C) circularity, D) type I collagen expression, and E) cadherin-11 expression reveals some contact-dependent differences between groups, but increased levels of activation in the M2 macrophage co-culture group even when fibroblasts did not directly contact macrophages. *N* = 5 hydrogels per group. Statistical analyses performed via two-way ANOVA with Tukey's HSD post-hoc testing. ****: $P < 0.0001$, **: $P < 0.01$, *: $P < 0.05$.

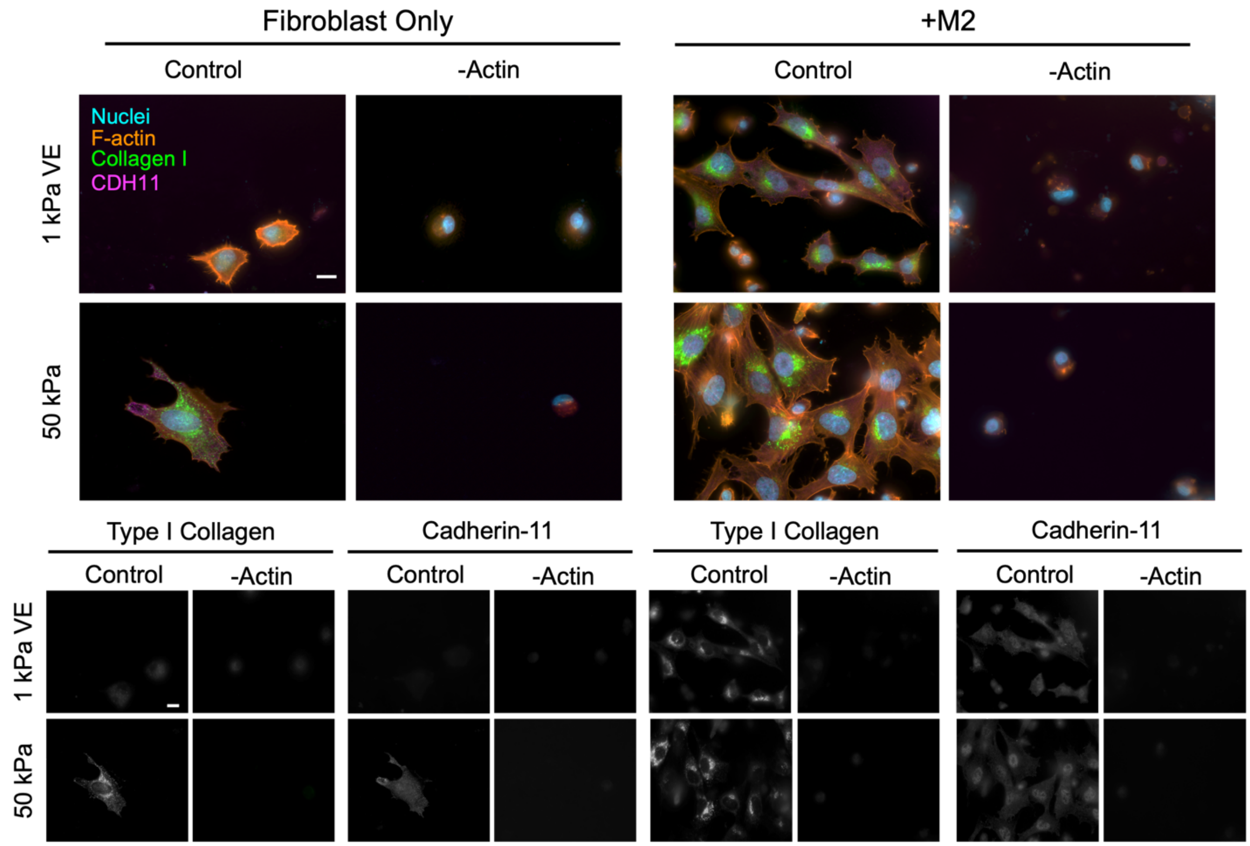

**Figure S5:** Inhibiting actin polymerization reduces fibroblast activation in all culture groups. A) Representative and single channel (type I collagen and CDH11 expression) images of fibroblasts cultured alone or with M2 macrophages on 1 kPa VE or 50 kPa hydrogels with or without actin polymerization inhibitor. Scale bars = 20 μm. Images shown from the no inhibitor control groups are reproduced for all inhibitor experiment figures (Fig. 7, S5-S8).

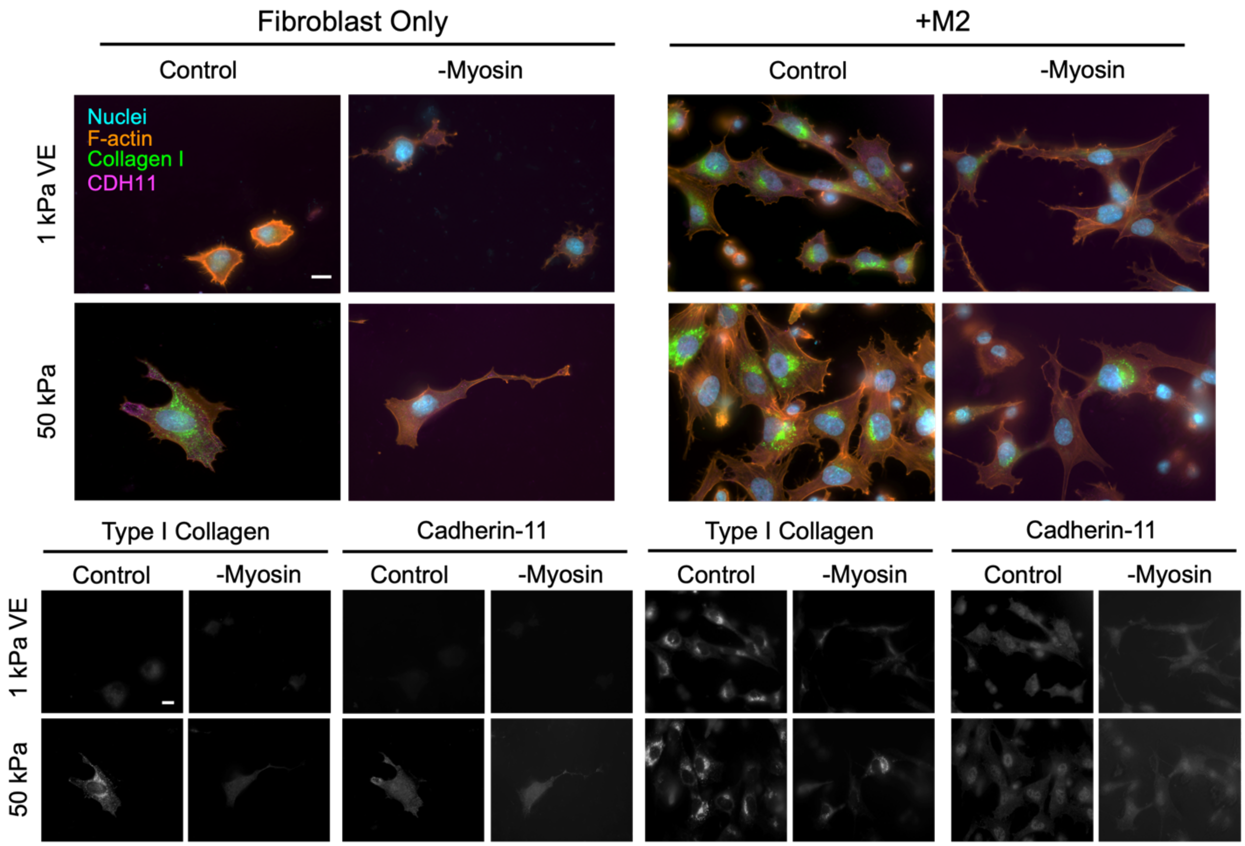

**Figure S6:** Inhibiting myosin II influences fibroblast activation by promoting decreased circularity and type I collagen expression. A) Representative and single channel (type I collagen and CDH11 expression) images of fibroblasts cultured alone or with M2 macrophages on 1 kPa VE or 50 kPa hydrogels with or without myosin II inhibitor. Scale bars = 20 μm. Images shown from the no inhibitor control groups are reproduced for all inhibitor experiment figures (Fig. 7, S5-S8).

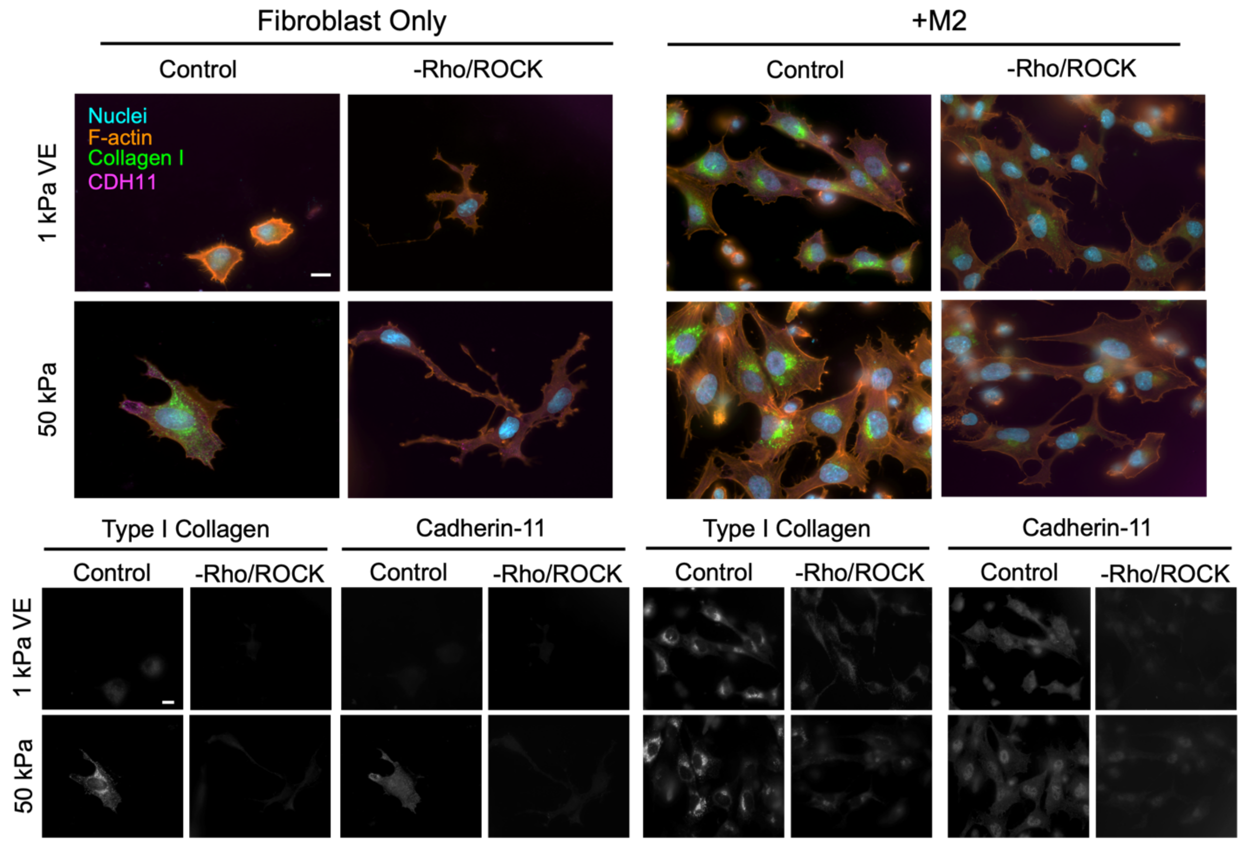

**Figure S7:** Inhibiting Rho kinase signaling influences fibroblast activation by reducing circularity, type I collagen expression in 50 kPa hydrogels, and CDH11 expression in co-culture groups. A) Representative and single channel (type I collagen and CDH11 expression) images of fibroblasts cultured alone or with M2 macrophages on 1 kPa VE or 50 kPa hydrogels with or without Rho kinase inhibitor. Scale bars = 20 μm. Images shown from the no inhibitor control groups are reproduced for all inhibitor experiment figures (Fig. 7, S5-S8).

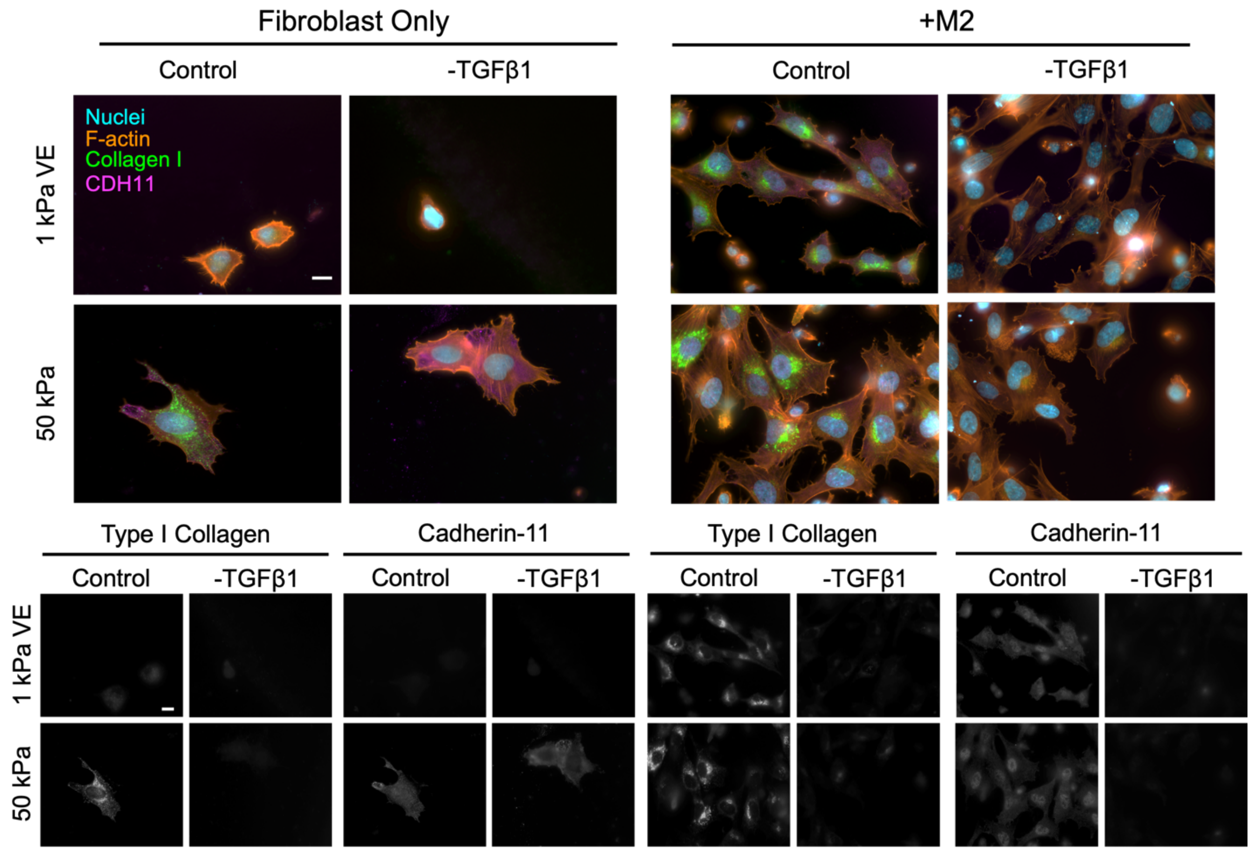

**Figure S8:** Inhibiting TGFβ1 signaling influences fibroblast activation by decreasing fibroblast spread area, type I collagen expression, and CDH11 expression in co-culture groups. A) Representative and single channel (type I collagen and CDH11 expression) images of fibroblasts cultured alone or with M2 macrophages on 1 kPa VE or 50 kPa hydrogels with or without TGFβ1 inhibitor. Scale bars = 20 µm. Images shown from the no inhibitor control groups are reproduced for all inhibitor experiment figures (Fig. 7, S5-S8).

### A Intragroup Comparison

#### Area

**1 kPa VE Fibroblast Only**

| | Control | TGFβ1 | Myosin II | Actin | Rho/ROCK | IL6 |
|---|---|---|---|---|---|---|
| Control | | 0.8509 | 0.2771 | <0.0001 | 0.104 | 0.9996 |
| TGFβ1 | | | 0.0281 | 0.0004 | 0.0079 | 0.9538 |
| Myosin II | | | | <0.0001 | 0.9936 | 0.1638 |
| Actin | | | | | <0.0001 | <0.0001 |
| Rho/ROCK | | | | | | 0.0554 |
| IL6 | | | | | | |

**50 kPa Fibroblast Only**

| | Control | TGFβ1 | Myosin II | Actin | Rho/ROCK | IL6 |
|---|---|---|---|---|---|---|
| Control | | <0.0001 | 0.385 | <0.0001 | 0.8404 | 0.806 |
| TGFβ1 | | | 0.0078 | 0.0001 | 0.0011 | <0.0001 |
| Myosin II | | | | <0.0001 | 0.9664 | 0.0371 |
| Actin | | | | | <0.0001 | <0.0001 |
| Rho/ROCK | | | | | | 0.1833 |
| IL6 | | | | | | |

**1 kPa VE + M2 Macrophages**

| | Control | TGFβ1 | Myosin II | Actin | Rho/ROCK | IL6 |
|---|---|---|---|---|---|---|
| Control | | 0.0027 | >0.9999 | <0.0001 | 0.6871 | 0.0001 |
| TGFβ1 | | | 0.0032 | <0.0001 | 0.0785 | 0.8434 |
| Myosin II | | | | <0.0001 | 0.728 | 0.0002 |
| Actin | | | | | <0.0001 | <0.0001 |
| Rho/ROCK | | | | | | 0.0054 |
| IL6 | | | | | | |

**50 kPa + M2 Macrophages**

| | Control | TGFβ1 | Myosin II | Actin | Rho/ROCK | IL6 |
|---|---|---|---|---|---|---|
| Control | | 0.0078 | 0.7988 | <0.0001 | 0.0079 | <0.0001 |
| TGFβ1 | | | 0.1265 | <0.0001 | >0.9999 | 0.0779 |
| Myosin II | | | | <0.0001 | 0.1278 | 0.0001 |
| Actin | | | | | <0.0001 | <0.0001 |
| Rho/ROCK | | | | | | 0.077 |
| IL6 | | | | | | |

#### CSI

**1 kPa VE Fibroblast Only**

| | Control | TGFβ1 | Myosin II | Actin | Rho/ROCK | IL6 |
|---|---|---|---|---|---|---|
| Control | | 0.0049 | <0.0001 | 0.3392 | <0.0001 | 0.9982 |
| TGFβ1 | | | <0.0001 | <0.0001 | <0.0001 | 0.0018 |
| Myosin II | | | | <0.0001 | 0.9955 | <0.0001 |
| Actin | | | | | <0.0001 | 0.5774 |
| Rho/ROCK | | | | | | <0.0001 |
| IL6 | | | | | | |

**50 kPa Fibroblast Only**

| | Control | TGFβ1 | Myosin II | Actin | Rho/ROCK | IL6 |
|---|---|---|---|---|---|---|
| Control | | 0.9926 | 0.0175 | <0.0001 | 0.0015 | 0.5007 |
| TGFβ1 | | | 0.061 | <0.0001 | 0.006 | 0.2192 |
| Myosin II | | | | <0.0001 | 0.9087 | 0.0002 |
| Actin | | | | | <0.0001 | 0.0039 |
| Rho/ROCK | | | | | | <0.0001 |
| IL6 | | | | | | |

**1 kPa VE + M2 Macrophages**

| | Control | TGFβ1 | Myosin II | Actin | Rho/ROCK | IL6 |
|---|---|---|---|---|---|---|
| Control | | 0.977 | 0.0124 | <0.0001 | 0.0832 | 0.0132 |
| TGFβ1 | | | 0.0022 | 0.0003 | 0.017 | 0.0664 |
| Myosin II | | | | <0.0001 | 0.9523 | <0.0001 |
| Actin | | | | | <0.0001 | 0.2468 |
| Rho/ROCK | | | | | | <0.0001 |
| IL6 | | | | | | |

**50 kPa + M2 Macrophages**

| | Control | TGFβ1 | Myosin II | Actin | Rho/ROCK | IL6 |
|---|---|---|---|---|---|---|
| Control | | 0.7099 | 0.986 | <0.0001 | >0.9999 | 0.3659 |
| TGFβ1 | | | 0.3326 | <0.0001 | 0.7276 | 0.9914 |
| Myosin II | | | | <0.0001 | 0.9829 | 0.1219 |
| Actin | | | | | <0.0001 | <0.0001 |
| Rho/ROCK | | | | | | 0.3821 |
| IL6 | | | | | | |

| |
|---|
| ns |
| * |
| ** |
| *** |
| **** |

#### Collagen

**1 kPa VE Fibroblast Only**

| | Control | TGFβ1 | Myosin II | Actin | Rho/ROCK | IL6 |
|---|---|---|---|---|---|---|
| Control | | 0.5411 | 0.8763 | 0.4945 | 0.466 | 0.7846 |
| TGFβ1 | | | 0.9896 | >0.9999 | >0.9999 | 0.0607 |
| Myosin II | | | | 0.9823 | 0.9761 | 0.1973 |
| Actin | | | | | >0.9999 | 0.0515 |
| Rho/ROCK | | | | | | 0.0464 |
| IL6 | | | | | | |

**50 kPa Fibroblast Only**

| | Control | TGFβ1 | Myosin II | Actin | Rho/ROCK | IL6 |
|---|---|---|---|---|---|---|
| Control | | <0.0001 | 0.0004 | <0.0001 | <0.0001 | 0.9993 |
| TGFβ1 | | | 0.001 | 0.9831 | 0.2165 | <0.0001 |
| Myosin II | | | | 0.0002 | 0.1946 | 0.001 |
| Actin | | | | | 0.0586 | <0.0001 |
| Rho/ROCK | | | | | | <0.0001 |
| IL6 | | | | | | |

**1 kPa VE +M2 Macrophages**

| | Control | TGFβ1 | Myosin II | Actin | Rho/ROCK | IL6 |
|---|---|---|---|---|---|---|
| Control | | <0.0001 | 0.6949 | <0.0001 | 0.9968 | 0.0003 |
| TGFβ1 | | | 0.0001 | 0.9906 | <0.0001 | 0.9581 |
| Myosin II | | | | <0.0001 | 0.4129 | 0.0012 |
| Actin | | | | | <0.0001 | 0.7116 |
| Rho/ROCK | | | | | | <0.0001 |
| IL6 | | | | | | |

**50 kPa +M2 Macrophages**

| | Control | TGFβ1 | Myosin II | Actin | Rho/ROCK | IL6 |
|---|---|---|---|---|---|---|
| Control | | <0.0001 | 0.0001 | <0.0001 | <0.0001 | <0.0001 |
| TGFβ1 | | | <0.0001 | 0.9998 | <0.0001 | >0.9999 |
| Actin | | | | <0.0001 | 0.9973 | <0.0001 |
| Myosin II | | | | | <0.0001 | 0.9998 |
| Rho/ROCK | | | | | | <0.0001 |
| IL6 | | | | | | |

#### CDH11

**1 kPa VE Fibroblast Only**

| | Control | TGFβ1 | Myosin II | Actin | Rho/ROCK | IL6 |
|---|---|---|---|---|---|---|
| Control | | 0.7416 | 0.2496 | 0.6923 | >0.9999 | 0.993 |
| TGFβ1 | | | 0.9467 | 0.0861 | 0.8495 | 0.4097 |
| Myosin II | | | | 0.0122 | 0.3468 | 0.0895 |
| Actin | | | | | 0.5637 | 0.9419 |
| Rho/ROCK | | | | | | 0.9707 |
| IL6 | | | | | | |

**50 kPa Fibroblast Only**

| | Control | TGFβ1 | Myosin II | Actin | Rho/ROCK | IL6 |
|---|---|---|---|---|---|---|
| Control | | 0.8952 | >0.9999 | 0.0032 | 0.9739 | 0.9988 |
| TGFβ1 | | | 0.9516 | 0.0002 | 0.9996 | 0.7057 |
| Myosin II | | | | 0.002 | 0.9929 | 0.9919 |
| Actin | | | | | 0.0005 | 0.0081 |
| Rho/ROCK | | | | | | 0.8644 |
| IL6 | | | | | | |

**1 kPa VE + M2 Macrophages**

| | Control | TGFβ1 | Myosin II | Actin | Rho/ROCK | IL6 |
|---|---|---|---|---|---|---|
| Control | | <0.0001 | 0.9029 | <0.0001 | <0.0001 | <0.0001 |
| TGFβ1 | | | <0.0001 | 0.0695 | 0.4108 | 0.8596 |
| Myosin II | | | | <0.0001 | <0.0001 | <0.0001 |
| Actin | | | | | 0.0007 | 0.0052 |
| Rho/ROCK | | | | | | 0.9671 |
| IL6 | | | | | | |

**50 kPa + M2 Macrophages**

| | Control | TGFβ1 | Myosin II | Actin | Rho/ROCK | IL6 |
|---|---|---|---|---|---|---|
| Control | | 0.0078 | 0.989 | <0.0001 | 0.0005 | <0.0001 |
| TGFβ1 | | | 0.0325 | 0.0112 | 0.8648 | 0.1114 |
| Myosin II | | | | <0.0001 | 0.0022 | <0.0001 |
| Actin | | | | | 0.1304 | 0.8971 |
| Rho/ROCK | | | | | | 0.6217 |
| IL6 | | | | | | |

### B Intergroup Comparison

**Area — Fibroblast Only v. +M2**

| | 1 kPa VE | 50 kPa |
|---|---|---|
| Control | <0.0001 | 0.8085 |
| TGFβ1 | 0.0013 | 0.0364 |
| Myosin II | 0.0001 | 0.351 |
| Actin | 0.5537 | 0.3248 |
| Rho/ROCK | 0.0145 | 0.0209 |
| IL6 | 0.1175 | <0.0001 |

**CSI — Fibroblast Only v. +M2**

| | 1 kPa VE | 50 kPa |
|---|---|---|
| Control | <0.0001 | 0.6782 |
| TGFβ1 | 0.2718 | 0.1292 |
| Myosin II | 0.1176 | 0.0199 |
| Actin | 0.0505 | 0.0159 |
| Rho/ROCK | 0.0593 | 0.0003 |
| IL6 | 0.0132 | 0.8568 |

**Collagen — Fibroblast Only v. +M2**

| | 1 kPa VE | 50 kPa |
|---|---|---|
| Control | <0.0001 | 0.1859 |
| TGFβ1 | 0.6111 | 0.5381 |
| Myosin II | 0.0002 | 0.364 |
| Actin | 0.3129 | 0.8418 |
| Rho/ROCK | <0.0001 | 0.0083 |
| IL6 | 0.013 | <0.0001 |

**CDH11 — Fibroblast Only v. +M2**

| | 1 kPa VE | 50 kPa |
|---|---|---|
| Control | <0.0001 | 0.0158 |
| TGFβ1 | 0.565 | 0.0269 |
| Myosin II | <0.0001 | 0.0853 |
| Actin | 0.4984 | 0.4403 |
| Rho/ROCK | 0.0194 | 0.004 |
| IL6 | 0.0203 | 0.0014 |

**Figure S9:** Tables of statistical comparisons from inhibitor data shown in Figures 6 and 7. A) Intragroup comparisons (within the same stiffness and culture type) of cell spread area, cell shape index, type I collagen, and cadherin-11 expression. B) Intergroup comparisons (comparing fibroblast-only data with M2 macrophage co-culture) of cell spread area, cell shape index, type I collagen, and cadherin-11 expression.

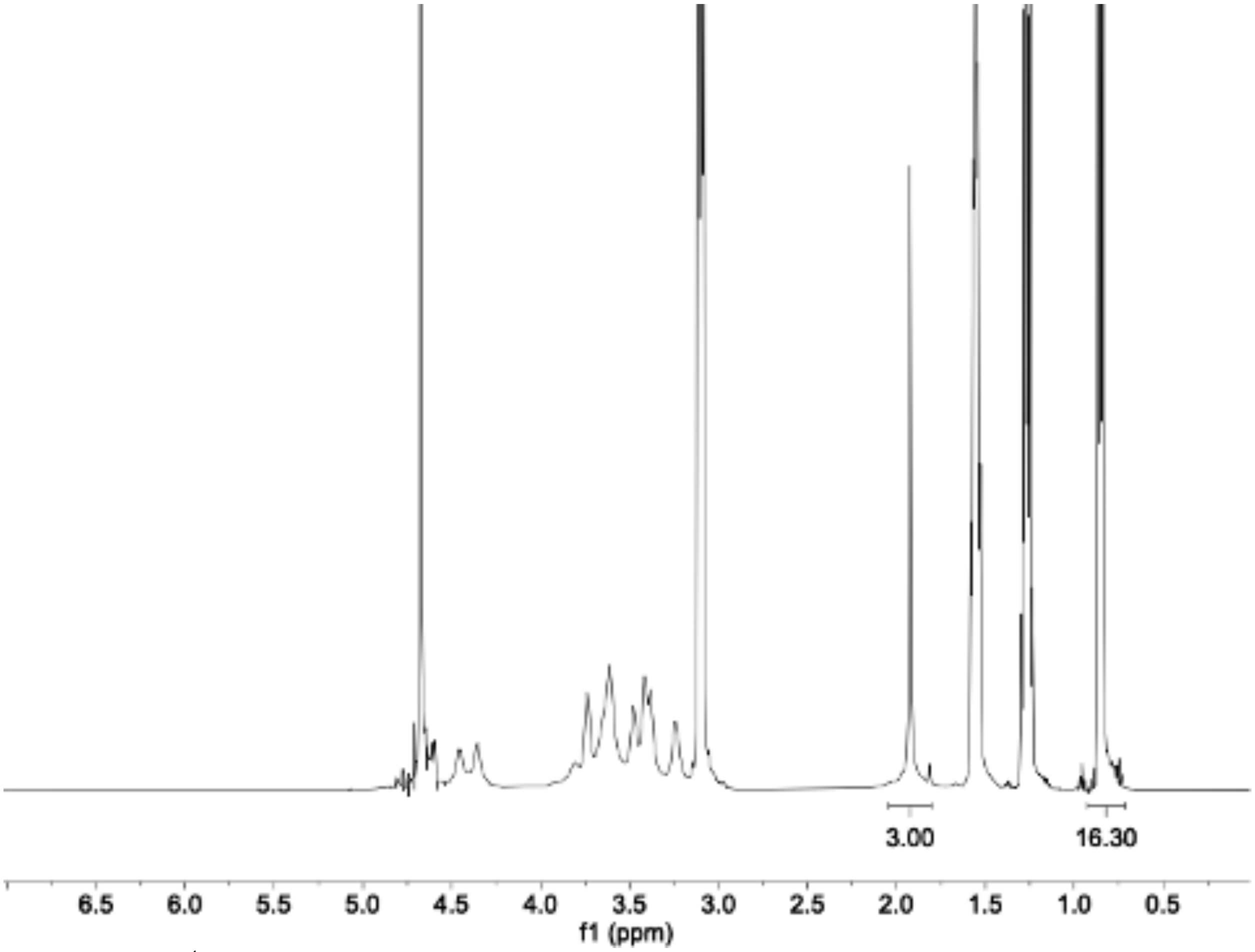

**Figure S10:** $^{1}$H NMR spectra of tetrabutyl ammonium salt of hyaluronic acid (HA-TBA). Modification of HA with TBA salt is determined by the integration of the TBA methyl groups relative to the *N*-acetyl group of HA.

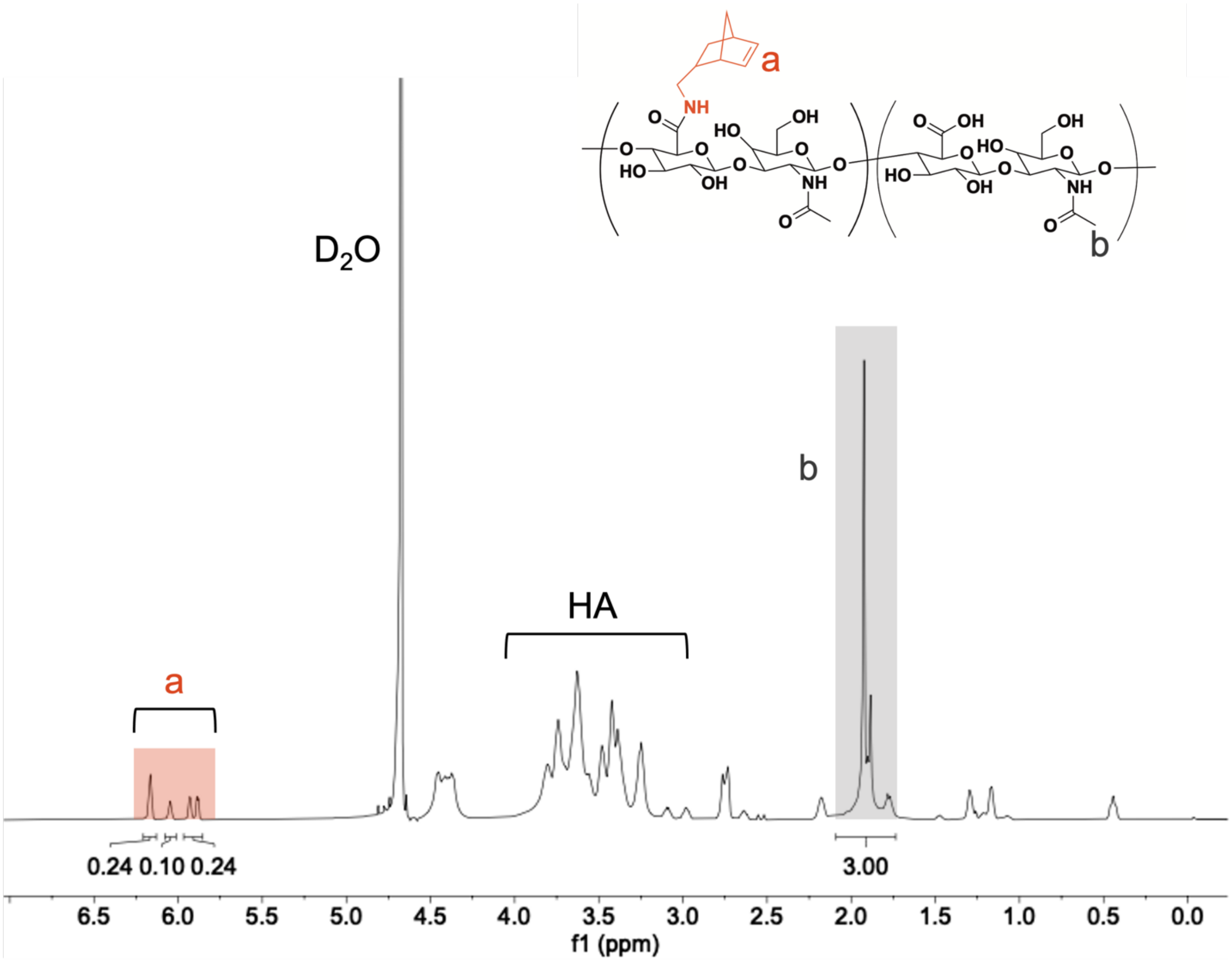

**Figure S11:** $^1$H NMR spectra of norbornene-modified hyaluronic acid (NorHA). Modification of HA with pendant norbornenes was determined to be 29%, indicated by the integration of peaks at $\delta$ = 5.75, 6.05, and 6.2 ppm (2H, ‘a’) normalized to the *N*-acetyl on HA (3H, ‘b’).

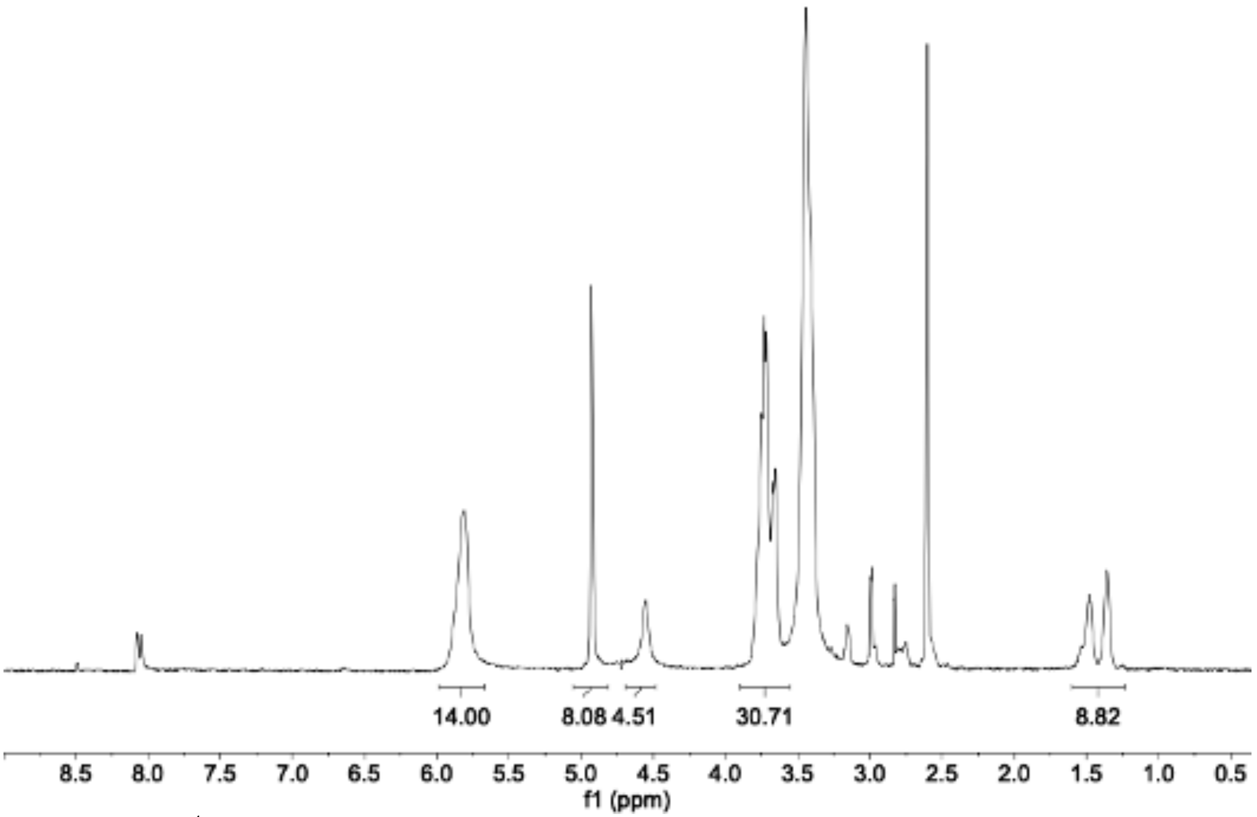

**Figure S12:** $^1$H NMR spectra of 6-(6-aminohexyl)amino-6-deoxy-β-cyclodextrin (CD-HDA). Modification of β-CD with HDA was determined to be 73%, indicated by the integration of peaks at $\delta$ = 1.14-1.6 ppm (12H).

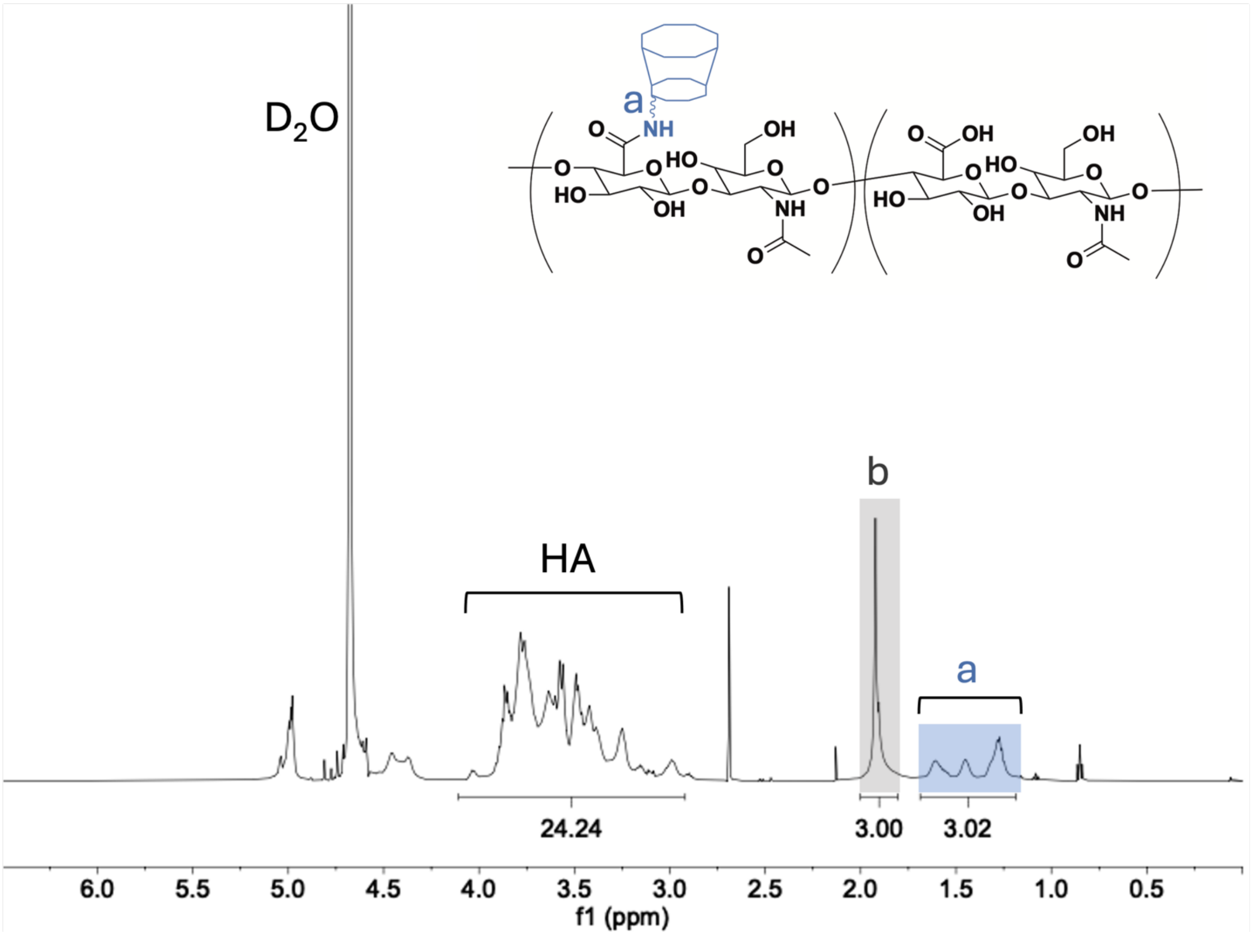

**Figure S13:** [1]H NMR spectra of β-cyclodextrin modified hyaluronic acid (CDHA). Modification of HA with pendant cyclodextrins (25%) was determined by integration of hexane linker peaks at $\delta$ = 1.2-1.7 ppm (12H, a) relative to the *N*-acetyl group of HA (3H, b).
